# Cellular transcriptomics reveals evolutionary adaptation and rumination of vertebrate stomachs

**DOI:** 10.64898/2026.03.09.710455

**Authors:** Qi-Xuan Huang, Song-Song Xu, Shan-Gang Jia, Wen-Tian Wei, Ya-Hui Zhang, Jing-Xuan Zhou, Jia-Hui Huang, Meng-Hua Li

## Abstract

Stomach, an essential digestive organ in vertebrates, has undergone significant changes in its morphology, chamber numbers, and physiology, in response to dietary diversity. However, its cellular and molecular basis underlying evolutionary adaptation remains largely uncharacterized. Here we report a single-cell and spatial transcriptomic atlas of stomachs from 23 vertebrate species with diverse feeding habits (e.g., omnivores, carnivores, and herbivores) and gastric architectures, including monogastric (e.g., primates), two- (e.g., aves), three- (e.g., camel) and four-chambered stomachs (e.g., sheep). We reveal conservation and divergence in cell-type composition, developmental trajectory, evolutionary origins and forces, spatial distribution, gene regulatory network and effect, metabolic signature and disease susceptibility associated with the feeding habits and stomach chamber. The diversification of plant based diets, particularly the expansion of coarse fibrous plants that required enhanced mechanical processing, driving the rapid evolution of specific cells and associated genes. Ruminants have evolved chamber-specific cell types with enhanced expressions of relevant functional genes (e.g., *KRT6A* in spinous cells of forestomach, *LUC7L* in SMCs of abomasum, and *TSPYL4* in abomasal enteroendocrine cells), which were validated by fluorescence in situ hybridization and spatial transcriptomic profiling. The three cell-specific expressed genes showed significant effects on cell proliferation and migration by RNA interference to knock down their expressions. In particular, the knockdown of *LUC7L* in SMCs promotes a transition from a contractile to a synthetic phenotype, and *Luc7l* knock-out in mice delays gastric emptying and impairs gastric motility, demonstrating that this gene is essential for the coordinated multi-chambered gastric motility that underlies rumination in ruminants. These findings elucidate the cellular and molecular adaptations underlying the evolution of stomach coupled with the emergence of diet land plants. Our results provide potential cellular and gene targets for engineering monogastric animals to acquire functional rumination digestive capabilities as well as for medication of gastric mobility.

## INTRODUCTION

Stomach is a crucial digestive organ which first evolved around 350–450 million years ago (Mya) as the vertebrate ancestors developed jaws^1,2^. In response to the evolutionary pressure of obtaining diverse food resources and enhancing adaptability^3^, the vertebrate stomach has diversified, developing into different feeding habits (e.g., omnivores, carnivores, and herbivores) and accordingly specialized structures with a varied number of 1–4 chambers (e.g., a single chamber in primates and four chambers of rumen, reticulum, omasum, and abomasum in ruminants)^4,5^. However, the complex evolutionary adaptation for vertebrate stomachs driven by cellular heterogeneity and distinct single-cell transcriptomic profiles remain largely unknown^6,7^. Additionally, the vertebrate stomach is susceptible to a range of gastric diseases (e.g., peptic ulcer, gastritis, and gastric cancer)^8^ with a low risk of gastric disorders in herbivores^9^. Thus, exploration of the cross-species cellular and molecular differences among vertebrate stomachs will help to understand the evolution of stomachs and its underlying genetic mechanisms.

Recent advances in single-cell RNA sequencing (scRNA-seq) and spatial transcriptomics have enabled the construction of comprehensive cross-species cell atlas, providing single-cell resolution insights into organ evolution such as lung^10^, intestine^11^, kidney^12^ and liver^13^. Despite these, the cellular composition and gene expression landscapes of both monogastric and polygastric stomachs remain incompletely understood with only a limited number of studies mostly in human and model animal species^14,15^. To date, no large-scale in-depth scRNA-seq studies have systematically compared stomach cellular composition across multiples species of different classes, in particular among herbivores with different number of stomachs (e.g., one for horse, three for camel, and four for cattle and sheep).

Here, we present the largest single-cell transcriptional atlas of gastric tissues, spanning 23 vertebrate species of differentiated feeding habits and structure (Fig. 1a).

**Fig. 1.**
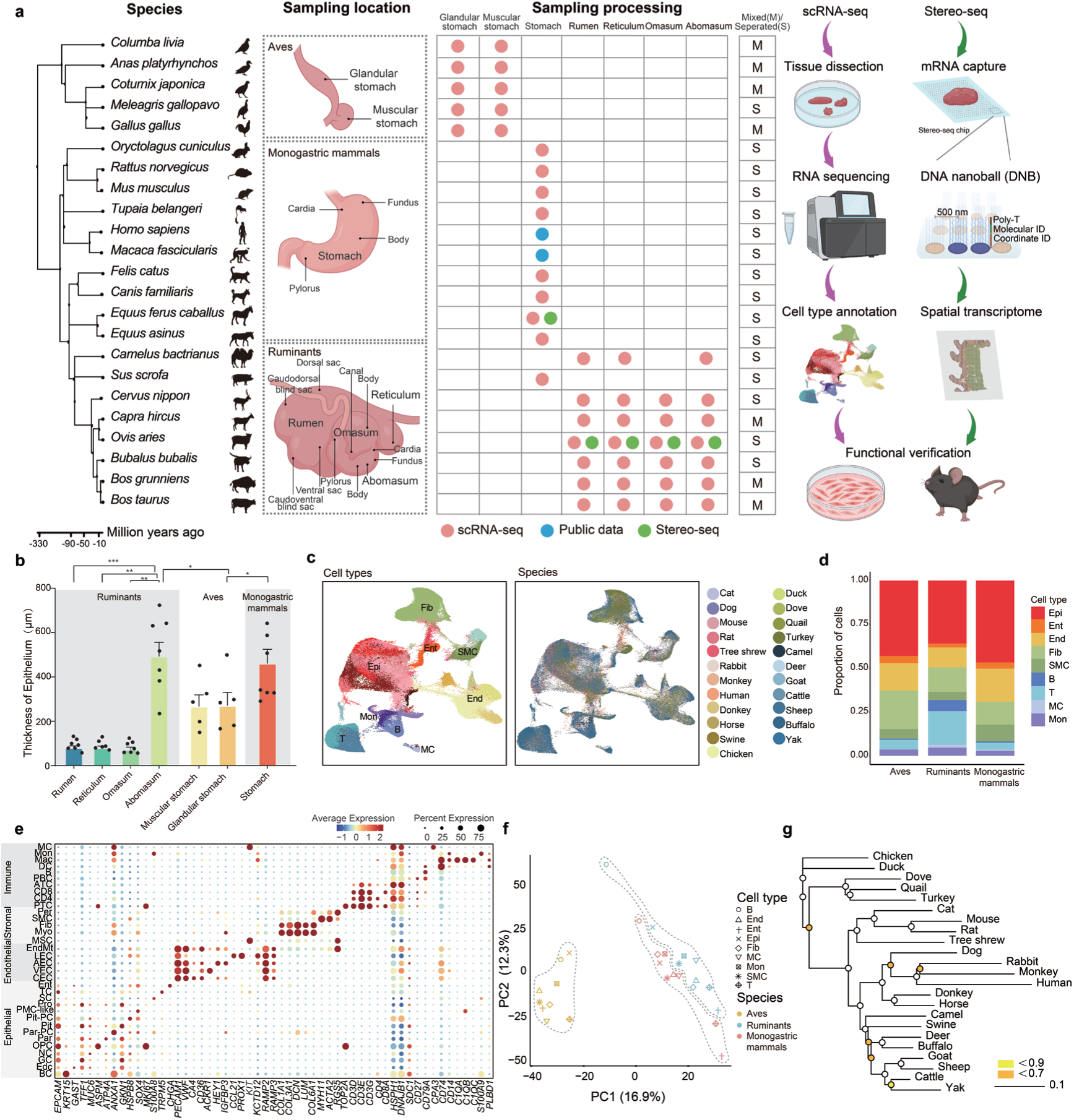
Cell atlas of stomach across 23 vertebrate species. a, Overview of study design. From left to right: the first column was the phylogenetic tree of 23 vertebrate species retrieved from the TimeTree; the second column was the multiple sampling locations from the seven stomachs (i.e., muscular and glandular stomachs in aves, stomach in monogastric mammals, and rumen, reticulum, omasum, and abomasum in ruminants); the third column was library construction strategies for scRNA-seq and Stereo-seq. “Mixed (M)” represents samples from multiple stomach chambers in a single species, and “Seperated (S)” indicates samples from individual stomach chambers in multichambered vertebrates; the fourth column was the experimental workflow for scRNA-seq, Stereo-seq, and functional verification. The images were created using BioRender. **b,** Epithelium thickness in rumen, reticulum, omasum, and abomasum of ruminants; muscular and glandular stomachs of aves; and stomach of monogastric mammals. Data are shown as mean ± SEM (standard error of the mean) with three independent experiments. *P* values are determined by Mann-Whitney U test, where ***, **, and * indicate *P* < 0.001, *P* < 0.01, and *P* < 0.05 respectively. **c,** Uniform manifold approximation and projection (UMAP) clustering of stomach cells by integrating scRNA-seq data from 23 species. (Left) Colors code different cell types. End, endothelial cells; Ent, enteroendocrine cells; Fib, fibroblast; Epi, epithelial cells; Mon, monocytes; B, B cells; T, T cells; SMC, smooth muscle cells; MC, mast cells. (Right) Same map as the left, but colors code 23 species. **d,** Cell type composition across monogastric mammals, aves, and ruminants. **e,** Expressions of gastric cell-type-specific canonical marker genes in immune, stromal, endothelial, and epithelial cells. **f,** Principal component analysis (PCA) of cell-type pseudo-bulks based on scRNA-seq. Monogastric mammals, aves, and ruminants are encircled by a dashed line. Each symbol represents an individual. **g,** Gene expression phylogeny based on pseudo-bulk transcriptomes for whole stomachs. Bootstrap values that were calculated by randomly sampling 3,682 1:1 orthologous genes with replacement 1,000 times are indicated by circles on the tree nodes, and colored in white for ≥0.9, yellow for <0.9, orange for <0.7.

Our findings uncovered conserved and divergent features in cell type distribution, evolutionary force, gene regulation, disease susceptibility, and functional specialization across these vertebrates. Additionally, we generated high-resolution spatial maps of cell types and gene expressions for the equine single-chambered stomach and the ovine four-chambered stomachs. The spatial organization of various cell types is highly structured, reflecting their adaptation to different digestive and absorptive functions. Notably, we identified and validated specifically-expressed genes (e.g., *KRT6A*, *TSPYL4*, and *LUC7L*) in certain cell types (e.g., spinous, enteroendocrine, smooth muscle cells) accounting for the rumination of ruminants. Our results provided critical insights into the cellular and molecular heterogeneity of gastric tissues associated with the evolutionary diversification of diet plant species, and advanced our understanding of evolutionary adaptation in the vertebrate digestive systems, in particular the rumination of ruminants.

## RESULTS

### A comprehensive single-cell transcriptome atlas of stomachs

To investigate the origin and evolution of stomachs, we first compared the stomach structure in 21 representative monogastric (nine species) and polygastric (twelve species) vertebrate species (Fig. 1a), including one-chambered mammal species (e.g., mouse, rat, tree shrew, cat, dog, horse, donkey, swine, and rabbit), two-chambered avian species (e.g., chicken, turkey, quail, duck, and dove), three-chambered ruminant species (e.g., camel), and four-chambered ruminant species (e.g., goat, sheep, yak, buffalo, cattle, and deer). Hematoxylin-eosin (HE) staining revealed histological differences among the seven stomachs of monogastric mammals, aves, and ruminants (Extended Data Fig. 1). For example, thickness of epithelium in the abomasum of ruminants and the stomach of monogastric mammals is significantly (*P* < 0.05, Mann-Whitney U test) higher than the glandular stomach of aves and the forestomaches (i.e., rumen, reticulum, and omasum) of ruminants (Fig. 1b). This could be explained by functions of the “true stomachs”, whose gastric glands secrete digestive enzymes, while its epithelial cells absorb nutrients^16^.

Furthermore, we generated scRNA-seq data for stomachs from these 21 species with omnivorous, carnivorous, and herbivorous diets. A total of 38 scRNA-seq libraries were constructed, generating 357,273,141–1,076,490,138 reads per library (Supplementary Table 1). Additionally, we collected stomach scRNA-seq datasets of monogastric human and monkey from public databases, expanding the analysis to encompass a total of 23 vertebrate species. Following stringent quality control, a total of 337,170 high-quality cells were obtained across 23 species, with an average of 14,660 cells per species. To harmonize these extensive datasets, we evaluated several widely used data integration tools and identified canonical-correlation analysis (CCA) as the top performer for effectively removing batch effects (Extended Data Fig. 2a, b). Uniform manifold approximation and projection (UMAP) embedding revealed a clear separation of cell types and effective integration across datasets (Fig. 1c). Next, we performed hierarchical unsupervised cell clustering and annotated nine broad cell types (i.e., epithelial, fibroblast, T cells, B cells, smooth muscle cells, endothelial, monocytes, mast cells, and enteroendocrine cells) using canonical markers and a host of new marker genes (Fig. 1c and Supplementary Table 2). The composition of some cell types exhibited significant variations among the monogastric mammals, aves, and ruminants (*P* < 2.2 × 10^−16^, Chi-squared test), and the abundance of immune cells (such as T and B cells) was substantially increased in ruminants (Fig. 1d and Extended Data Fig. 2c). Further, the nine broad cell types were divided into 34 cell subtypes. For example, the epithelial cells comprised 14 subtypes such as basal, granular, spinous, and pit cells. Endothelial cells encompassed artery, vein, capillary, and lymphatic subtypes, while the immune cells included 10 diverse cell types like mast cells, B cells, monocytes, and macrophages (Fig. 1c, e and Supplementary Table 3). High score of area under the receiver operator characteristic curve (AUROC) indicates the accuracy of annotation and assignment of cell types and subtypes (Extended Data Fig. 2d).

To explore the cross-species similarity among cell types, principal component analysis (PCA) was performed based on pseudo-bulk transcriptomes. Aves were clustered into a group away from the monogastric mammals and ruminants by PC1, agreeing with their phylogenetic framework (Fig. 1f). The divergence between ruminants and monogastric mammals could also be observed in the PCA plot (Fig. 1f). To further investigate gene expression evolutionary rate across the cell types, we built a gene expression phylogenetic tree based on pseudo-bulk transcriptomes (Fig. 1g). The evolutionary relationships of all the 23 species were in accordance with the clustering tree of the whole cell types (Fig. 1g), suggesting that gene expression changes experienced differential accumulation over evolutionary time in animal lineages and species. Collectively, these results provide a single-cell atlas of stomach across 23 vertebrate species, encompassing over 330 million years of evolutionary divergence.

### Differential evolutionary forces and novel candidate disease genes

To trace the cellular source of stomach’s evolution, we constructed expression trees for individual cell types (Extended Data Fig. 3a). The total branch lengths of these trees differ considerably among cell types, indicating the variable extent of evolutionary expression change (Fig. 2a). We observed substantially higher rate of expression evolution in the immune cells (e.g., B and mast cells). Additionally, the relationship between Spearman’s *ρ* and divergence time further confirmed the rapid expression evolution of immune cells across species and the increase of gene expression divergence with evolutionary time (Fig. 2b). To characterize the evolutionary forces of each cell type, we calculated the average normalized ratio of non-synonymous to synonymous substitutions (dN/dS) across the 23 species^17^. The immune cells (e.g., B and mast cells) and enteroendocrine cells showed higher dN/dS ratios than the other cell types, implying less constraint on the protein-coding genes expressed in these cells (Fig. 2c). Notably, enteroendocrine-enriched genes with high dN/dS ratios in ruminants are involved in cell adhesion and glycerolipid metabolism (e.g., *CD47, SERPINF2, SKAP1, XBP1*, and *GPCPD1*) (Fig. 2e and Supplementary Table 4). We also found that the dN/dS ratios of most cell types (except for endothelial and mast cells) are higher in ruminants (Fig. 2d) than those in monogastric mammals and aves, suggesting a higher proportion of genes under positive selection.

**Fig. 2.**
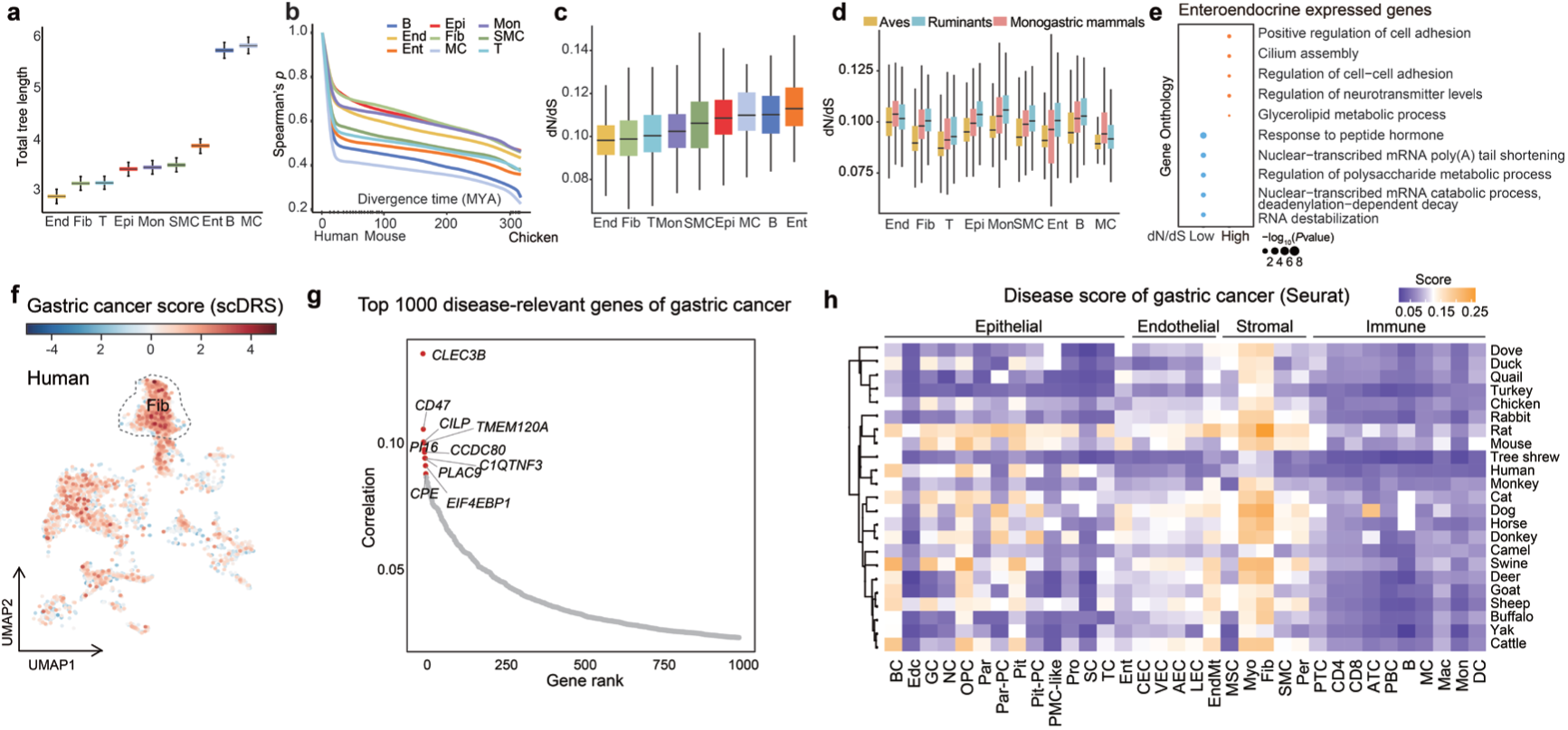
Gene expression divergence, evolutionary forces, and gastric cancer relevance across 23 species. **a**, Total branch lengths of expression trees among nine broad stomach cell types across 23 vertebrate species. Box plots show the median (central value); upper and lower quartile (box limits) and 95% confidence intervals (whiskers) for 1,000 bootstrap replicates. **b,** Plot indicating evolutionary distance (x-axis) and transcriptional correlation (y-axis) for each cell type (indicated by colors) between humans and the other species. **c,d,** Mean normalized ratio of nonsynonymous (dN) over synonymous (dS) nucleotide substitutions of expressed genes is plotted across nine broad cell types in all species (**c**), and compared among monogastric mammals, aves, and ruminants (**d**). **e,** Gene ontology enrichment analysis for enteroendocrine-cell-enriched genes with high dN/dS scores (> 0.8). **f,** Association of human gastric cells with gastric cancer. Significantly associated cells (FDR < 0.1) are denoted in red, with shades of red denoting scDRS disease scores, and the other cells are denoted in gray. **g,** The top 1,000 disease-relevant genes of gastric cancer risk in human. The correlation coefficient for each gene was computed using scDRS. The top 10 genes most strongly associated with disease risk are highlighted in red and shown with gene symbols. **h,** The disease scores (Seurat) were calculated on *Addmodulescore* in Seurat based on the top 1,000 most relevant genes for gastric cancer (scDRS predicted), and plotted in a heatmap across the cell types (full cell names shown in Supplementary Table 17) in 23 vertebrate species.

Previous studies reported evolutionary links between diet and cancer vulnerability across species, and notably, ruminants are the least cancer-prone mammalian animals^9^. To understand the lower gastric cancer risk in ruminants, we applied scDRS to integrate scRNA-seq data with genome-wide association studies (GWAS) results for gastric cancer in humans^18–20^ (Fig. 2f). Gastric cancer showed strong associations (scDRS disease score difference >2) with the fibroblasts in humans (Fig. 2f). Further analysis revealed that a set of genes (e.g., *CLEC3B*, *CD47*, *CILP*, and *PI16*) showed strong correlations with gastric cancer (Fig. 2g). Among these, *CLEC3B* exhibited the highest correlation with gastric cancer risk (Fig. 2g) and was previously regarded as a potential tumor suppressor biomarker for cancer in humans^21^. Its elevated expression in ruminant endothelial cells (Extended Data Fig. 3c) may contribute to their reduced susceptibility to gastric cancer through its anti-angiogenic function^22,23^. Across the 23 species, fibroblasts showed the highest scores in gastric cancer (Fig. 2h). This indicated that fibroblasts might participate in the malignant development of gastric cancer^24^. Notably, the marker genes of fibroblasts (Supplementary Table 2), such as *CLEC3B*, *PI16*, *CCDC80*, *DCN*, *COL1A2*, *COL3A1*, and *COL12A1*, showed high scDRS scores for gastric cancer. Thus, our results suggested these marker genes of fibroblasts represented a novel set of candidate genes and biomarkers for gastric cancer (Supplementary Table 5).

### Conservation and divergence of gastric cell types and gene expressions across species

Based on single-cell expression data, MetaNeighbor analysis revealed a clear divergence of epithelial cells across the 23 species (AUROC < 0.75), in contrast to the high conservation observed in the endothelial and stromal cells (Fig. 3a). We uncovered a set of novel cell subtypes, e.g., a *COL4A6*^+^ fibroblast subtype in aves and a *CCR10*^+^ B memory cell subtype in ruminants (Extended Data Fig. 4a). *COL4A6*, a type IV collagen, is a major structural component of the basement membrane (BM), and could be involved in maintaining the integrity of the gastric mucosa^25^. CC Motif Chemokine Receptor 10 (*CCR10*) participates in the maintenance of immune homeostasis in the stomach by regulating the trafficking of B memory cells^26^. The analysis of marker genes for each cell type against the others showed the conservation for certain marker genes across the three groups of aves, ruminants, and monogastric mammals, but the majority of marker genes were uniquely enriched in a single group (Fig. 3b and Supplementary Table 6). The marker genes shared by aves, ruminants and monogastric mammals in epithelial (*n* = 51), B (*n* = 41), and mast (*n* = 45) cells were less than those in the other cell types (e.g., 552 and 251 marker genes for endothelial cells and SMCs, respectively), suggesting their divergence and significance (Fig. 3c and Extended Data Fig. 4b).

**Fig. 3.**
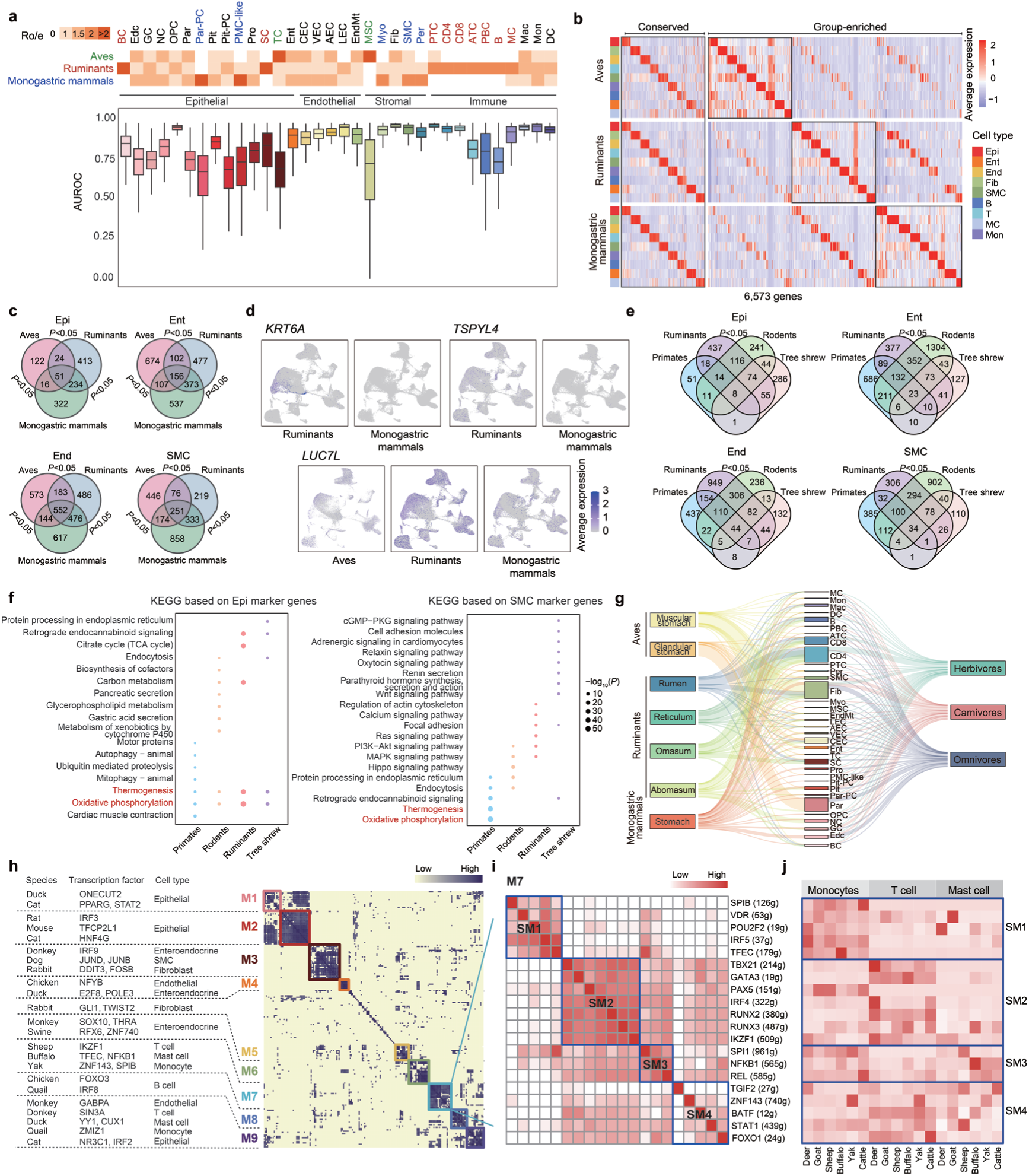
Conserved and divergent features of gastric cell types, genes, and regulon modules across 23 species. **a**, Preference of each cell type among monogastric mammals, aves, and ruminants, evaluated by the Ro/e index (ratio of observed to expected cell number, top) and AUROC (bottom). The scale of Ro/e index is placed on the top left corner, and the cell types that are uniquely present in one single animal group are highlighted in three colors, green for aves, red for ruminants, and blue for monogastric mammals. Box plot showing total AUROC scores from each cell subtype from endothelial, epithelial, stromal, and immune cells. Center line, box bounds, and whiskers represent mean, 25^th^ to 75^th^ percentile range, and minimum to maximum range, respectively. **b,** Heat map showing the expression of conserved (1554 genes) and group-enriched (5019 genes) marker genes for each broad cell type among monogastric mammals, aves, and ruminants. **c,** Venn diagrams showing the conserved and divergent marker genes for four broad cell types among monogastric mammals, aves, and ruminants. **d,** UMAP plots showing the expression distributions of *KRT6A*, *TSPYL4*, and *LUC7L* in the stomach cells of monogastric mammals, aves, and ruminants. **e,** Venn diagrams showing the marker genes of four broad cell types among primates, ruminants, rodents, and tree shrew. **f,** Enriched KEGG pathways for marker genes of epithelial and smooth muscle cells in primates, ruminants, rodents, and tree shrew. **g,** Sankey plot shows relationship of cell types among the stomach chambers of monogastric mammals, aves, and ruminants, and with three feeding habits (i.e., omnivores, carnivores, and herbivores). **h,** Regulon modules based on regulon connection specificity index (CSI) matrix, along with representative transcription factors and associated cell types. **i,** Zoomed-in view of module M7 identifies sub-modules SM1–SM4. **j,** Sub-modules SM1–SM4 in M7 are associated with distinct immune cell types and regulon activities. Epi, epithelial cells; Ent, enteroendocrine cells; End, endothelial cells; SMC, smooth muscle cells. The full names of cell types are shown in Supplementary Table 17.

Notably, we found that three genes *TSPYL4*, *KRT6A* and *LUC7L* were highly expressed in ruminants compared to aves and monogastric mammals (Fig. 3d, Extended Data Fig. 4d and Supplementary Table 7). *TSPYL4* is abundantly expressed in enteroendocrine cells, and is absent in the avian genomes (Extended Data Fig. 4d, e). This gene were previously reported to be involved in gastric epithelial repair and cell migration^27–29^, consistent with the results of our integrated transcriptomic and proteomic analyses and cellular functional assay (Extended Data Fig. 5). As a key marker of spinous cells, *KRT6A* is predominantly localized in the stratified squamous epithelium of the forestomach (i.e., rumen, reticulum, and omasum)^30^, where it enhances cytoskeletal stability to help spinous cells resist mechanical abrasion from coarse forage. Cellular functional assays indicated that *KRT6A* knockdown in rumen epithelial cells significantly impaired scratch wound healing and reduced cell proliferation, which might be mediated by the enriched pathways of G protein-coupled receptor signaling^31^, Wnt signaling pathway^32^, JAK-STAT^33^ and FoxO signaling pathway^34^ by transcriptomic analysis (Extended Data Fig. 6). *LUC7L* has been reported to play an important role in the regulation of muscle differentiation^35^. The functional role of *LUC7L* was further validated through cellular and *in vivo* experiments, as detailed in the results section below.

Ruminants generally exhibit higher efficiency in digesting cellulose and absorbing nutrients from fibrous plant materials compared to monogastric herbivores^36^. Accordingly, we conducted a comparative analysis between the stomach of monogastric herbivores (e.g., horses, donkeys, and rabbits) and the abomasum of ruminants (e.g., camels, deers, and buffalos). Our AUROC analysis revealed strong conservation of cell types, in particular endothelial cells (Extended Data Fig. 4c), across the six species. The endothelial cells are dominant in the stomach of monogastric herbivores (Extended Data Fig. 2c), and exhibited a higher proportion than those of ruminants. In contrast, epithelial cells in horses, donkeys, and rabbits are clustered away from the ones of ruminants’ abomasum, and exhibited the biggest divergence (Extended Data Fig. 4c). Epithelial cells were less abundant in monogastric herbivores than those in the other ruminants (Extended Data Fig. 2c). By the time digesta reaches the abomasum from the forestomach, it has already undergone substantial pre-digestion, facilitating efficient nutrient absorption by epithelial cells^5^. In fact, monogastric herbivores primarily rely on hindgut fermentation, where their stomachs play a minimal role in pre-digestion^5^, suggesting distinct digestive strategies between monogastric herbivores and ruminants.

Meanwhile, we compared marker genes in ruminants with those in primates and the model animals such as rodents (e.g., mice and rats) and tree shrew. We observed that ruminants shared a greater number of marker genes with rodents than those shared with primates and tree shrew (Fig. 3e and Extended Data Fig. 7a), suggesting their great conservation of gene regulation. Subsequent KEGG analysis of the cell type-specific marker genes revealed distinct pathway enrichment patterns across the four animal groups (Fig. 3f and Extended Data Fig. 7b). Across all the four animal groups, the oxidative phosphorylation and thermogenesis pathways were significantly enriched in epithelial cells (Fig. 3f and Extended Data Fig. 7b). In contrast, no enrichment of oxidative phosphorylation and thermogenesis pathways was observed in the other cell types like smooth muscle, endothelial, and B cells (Fig. 3f and Extended Data Fig. 7b).

### Diversity of cell types and cellular interactions associated with feeding habits and stomach chambers

We compared the cellular heterozygosity among the seven stomach samples from monogastric mammals (one), aves (two), and ruminants (four), and then among the three animal groups defined by the feeding habits (i.e., omnivores, carnivores, and herbivores). We observed enrichment of specific cell subtypes for the seven stomachs, such as fibroblasts for muscular stomach, parietal cells for glandular stomach, CD4^+^ T cells for rumen, reticulum, and omasum, neck cells for abomasum, and endocrine cells for monogastric stomach (Fig. 3g and Extended Data Fig. 6a). Also, immune-related cells are abundant in herbivores, such as CD4^+^ T cells, and the epithelial and stromal cells are increased in carnivores (e.g., pit cells, SMCs, and fibroblasts) and omnivores (e.g., parietal cells, endocrine cells, and fibroblasts) (Fig. 3g).

Also, we explored the conservation of ligand‒receptor pairs underlying the cell-cell interactions among the nine broad cell types, and identified three conserved ligand‒receptor pairs in monogastric mammals, 86 in aves, and 136 in ruminants (Supplementary Table 8). The ligand‒receptor pair NRP2-SEMA3C was commonly identified across the 23 species (Extended Data Fig. 8b), suggesting its conserved role in vascular angiogenesis^37,38^. In monogastric mammals, the conserved ligand‒receptor pairs, such as the VEGFA-FLT1 pair, are associated with vascular angiogenesis^39^, which are critical for the absorption and transport of nutrients. Meanwhile, the conserved ligand‒receptor pairs in aves, including FGF2-FGFR1 and WNT5A-ROR1, were enriched in cell proliferation and migration pathways^40,41^, whereas ruminants exhibited a pronounced immune regulatory network involving TNF-TNFRSF1A^42^ and TGFB1-TGFBR3^43^ (Extended Data Fig. 8c and Supplementary Table 8). We also discovered the functional divergence of ligand‒receptor pairs for the dietary adaptations (Extended Data Fig. 8d). The common enriched pathways for the ligand-receptor pairs were identified in both carnivores and herbivores, such as peptidyl−tyrosine phosphorylation, peptidyl−tyrosine modification, and positive regulation of kinase activity (Extended Data Fig. 8d). In contrast, the group-specific pathways were also detected, such as mesenchyme development and epithelial migration in herbivores, and neuron-related pathways in omnivores (Extended Data Fig. 8d). Together, we systematically revealed those highly conserved and lineage-specific cell-to-cell signaling within vertebrate stomachs associated with distinct feeding habits across species.

### Evolutionary conservation and divergence of gastric regulatory networks

To gain further insight into the combinatorial patterns of transcription factors (TFs) and gene expression levels, we calculated the atlas-wide similarity of the regulon activity scores (RAS) of every regulon pair by utilizing the Connection Specificity Index (CSI)^44^. A total of 155 regulons were identified and clustered into nine major modules (Fig. 3h). The module M7, which contains 20 regulons, presented the highest specificity score in ruminants. This module is pronounced in immune cell types, such as T cells, mast cells, and monocytes (Fig. 3h). Due to the complexity of ruminants’ immune system, we divided M7 into four smaller sub-modules (SM1–SM4) (Fig. 3i). Each sub-module is specifically related to distinct immune cell types (Fig. 3j). For example, SM1 is associated with monocytes, and the involved regulons including *SPIB*^45^ and *TFEC*^46^ regulate immune cells development, activation, and function. SM2 is mainly related to T cells, and contains many well-known regulators related to T cells, such as *IKZF1*^47^, *RUNX3*^48^, *GATA3*^49^ and *IRF4*^50^. Regulons in SM3 (such as *NFKB1*, *SPI1* and *REL*) are highly active in mast cells, regulating the activation, cytokine production, and survival of immune cells^51^. These results indicated that the ruminant-specific regulatory network (M7) is composed of specialized sub-modules which function cooperatively to coordinate specific pathways within the immune system.

We also identified four common TFs (i.e., ATF3, GABPA, NFKB1, and STAT1) across 23 species, suggesting their evolutionary conservation in vertebrate species^52–54^. We then constructed the TF regulatory networks for each species to identify their candidate target genes (Extended Data Fig. 8e). Specifically, the regulons of ATF3-DUSP1 and ATF3-EGR1 were identified across all the species. *DUSP1* plays an important role in the cellular response to environmental stress as well as in the negative regulation of cellular proliferation^55^, while *EGR1* activation is required for differentiation and mitogenesis^56^. This suggests that the regulons in stomachs are evolutionarily ancient and conserved in vertebrates.

### Cellular heterogeneity of ruminants’ chambers

Ruminants exhibited distinct cellular composition preferences compared to aves and monogastric mammals (Extended Data Fig. 2c). Further, we performed a comparative analysis of cell subtypes and gene expression patterns across the individual stomach chambers of ruminants like camels (lacking omasum^57^), deer, yak, and sheep, whose samples have been collected for each chamber separately (Extended Data Fig. 9a). We observed the conserved cellular composition of the abomasum across the four ruminant species (Extended Data Fig. 9a). Meanwhile, we identified biased distribution of cell types between forestomach (i.e., rumen, reticulum, and omasum) and abomasum (Fig. 4a and b). For example, forestomach epithelium consists of stratified squamous epithelial cells containing basal, pro_basal, granular, and spinous cells (Fig. 4b and Extended Data Fig. 9b, c), which together form a specialized structure with protective and absorptive functions, unlike the enzyme-secreting epithelium of the abomasum^4,58^. In contrast, the abomasum resembles the structure and function of the stomach of monogastric animals^59^, and harbors unique epithelial cell types which are absent in forestomach, such as enteroendocrine and pit cells (Fig. 4b and Extended Data Fig. 9b). The refined epithelium differentiation enables ruminants to efficiently utilize plant fiber resources^60^. Specifically, with fermentation occurs primarily in forestomach, while nutrient absorption is localized to abomasum, demonstrating the unique adaptability and high efficiency of the ruminant digestive system. Nevertheless, forestomach manifested species-specific cellular differences, for example, dominant B cells in camel reticulum, fibroblasts in deer rumen and reticulum, granular cells in buffalo, and CD4^+^ T cells in sheep (Extended Data Fig. 9a).

**Fig. 4.**
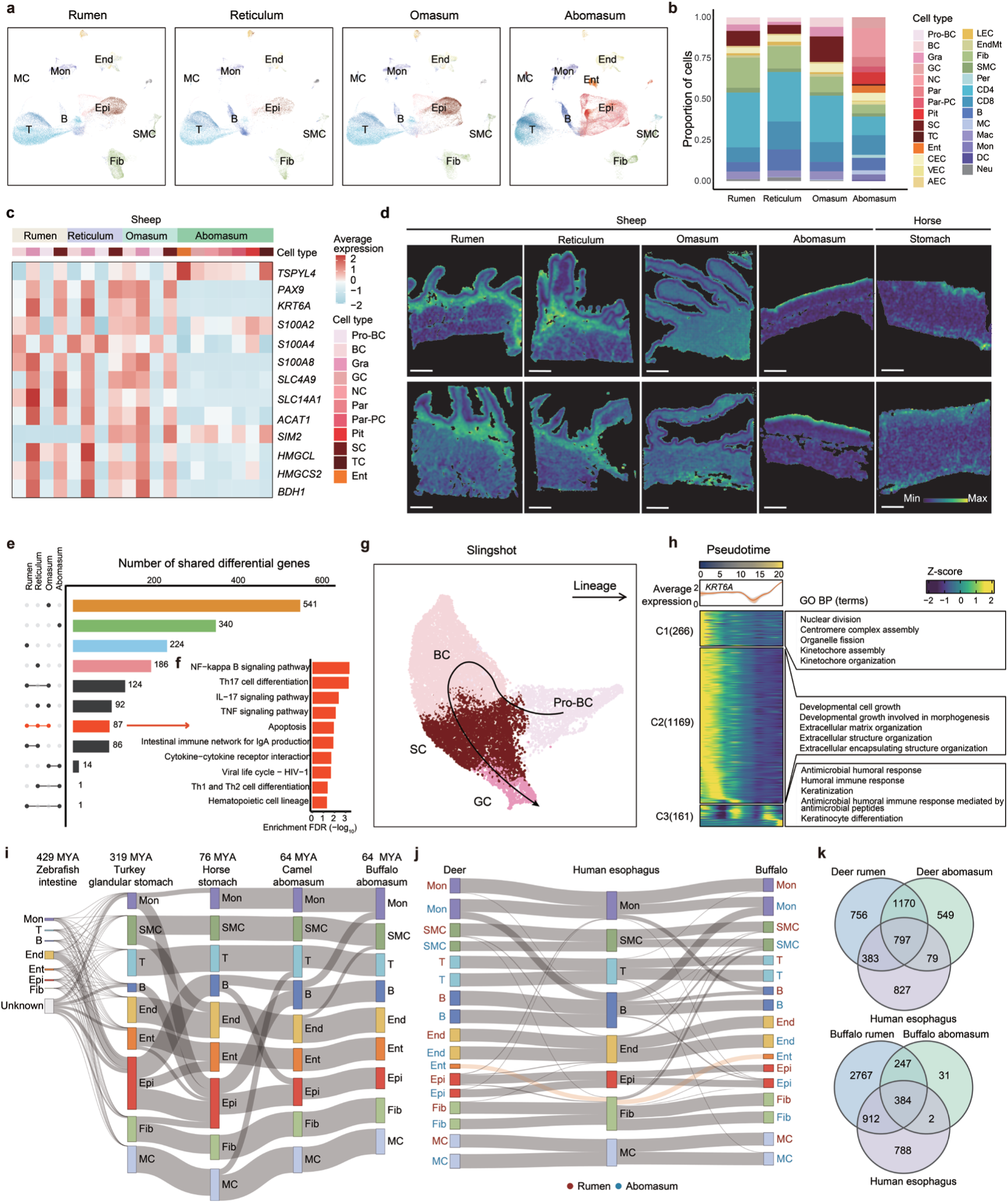
Functional specialization, differentiation trajectories, and evolutionary origins of the ruminant stomach chambers. a, UMAP visualization of cell type distribution of stomach chambers in ruminants (sheep, deer, buffalo, and camel). **b,** Cell-type composition among the ruminant stomach chambers. **c,** Heatmap showing expression pattern of the selected genes previously published in stomach studies in epithelial cell subtypes of the four sheep stomach chambers. **d,** Log-normalized spatial expression of the same selected candidate genes (**c**) across four stomach chambers of two sheep and two equine stomachs. The stomach mucosal (inner) surface faces upward in each panel. **e,** UpSet plot of the differentially expressed genes (DEGs) among the stomach chambers in ruminants (sheep, deer, buffalo, and camel) based on orthologous genes. Bars indicating DEGs uniquely found in each stomach chamber are colored. The bar for the DEGs shared by the rumen, reticulum, and omasum is colored in red and these DEGs are enriched in KEGG pathways (**f**). **g,** A UMAP plot showing the trajectory of epithelial cells from pro_basal cells (Pro-BC), basal cells (BC), spinous cells (SC), to granular cells (GC) in the forestomachs of ruminants (sheep, deer, buffalo, and camel). **h,** Heatmap showing dynamic expression of three clusters of genes (C1, C2, and C3) along pseudotime of epithelial cells in the forestomachs. Z scores indicate relative expression (yellow = high, blue = low). GO biological process (BP) terms associated with each cluster are displayed on the right, while the expression pattern of *KRT6A* is shown along the pseudotime trajectory at the top. **i, j,** Sankey plot showing the evolution of cell types from the glandular stomach of turkey, stomach of horse, abomasum of camel and buffalo, and intestine of zebrafish (**i**) and among the human esophagus, rumen, and abomasum in both buffalo and deer (**j**). **k,** Venn diagrams showing the expressed genes shared among the human esophagus, rumen, and abomasum in both deer (top) and buffalo (bottom). The full names of cell types are shown in Supplementary Table 17.

### Distinct gene expression and metabolic profiles among the chambers of ruminants

We examined single-cell gene expression patterns for each stomach chamber in ruminants to characterize their distinct patterns. We collected previously reported genes related to rumen physiological and rumination^4,61^, and found that these genes consistently exhibited elevated expression levels in epithelial cells in ruminants (Extended Data Fig. 9d). Most of these genes (e.g., *KRT6A* and *S100A8*) were highly expressed in forestomach, while *TSPYL4* showed high expression in the abomasum (Fig. 4c). We visualized their localization using spatial transcriptomics, and found their enrichment in the epithelium layer in the four stomach chambers of ruminants and the stomach of horses (Fig. 4d). Association of these genes with epithelial functions, as established in previous studies, is confirmed spatially by our data, thereby affirming its reliability. We found a total of 1,696 differentially expressed genes (DEGs) across the four chambers in a pseudo-bulk RNA-seq analysis based on our scRNA-seq data (Fig. 4e). Among them, 1,291 DEGs are chamber-specific and uniquely expressed in each of four stomach chambers, with the greatest number in omasum (Fig. 4e). These chamber-specific DEGs are strongly related to the unique pathways in the four chambers, and the neighbor chambers of rumen and reticulum share immune pathways, such as tumor necrosis factor (TNF) signaling pathway and Th17 cell differentiation (Fig. 4f and Extended Data Fig. 9e). The enriched DEGs of abomasum are associated with gastric acid secretion, which is also required in the stomach of monogastric mammals. The enriched DEGs of omasum are involved in metabolism, such as cysteine and methionine metabolism and carbon metabolism.

Due to few DEGs between abomasum and forestomach, we focused on forestomach and identified 87 common DEGs, which are enriched in immune-related pathways (Fig. 4f). This could be explained by the fact that the immune pathways play a role in balancing microbial symbiosis and precise defense.

To systematically evaluate metabolism differences at the single-cell level among the four stomach chambers, we employed an integrated algorithm named single-cell flux estimation analysis (scFEA)^62^, for the nine cell types of scRNA-seq. The epithelial cells of the rumen exhibited an increase in the fluxes of the fatty acid metabolism, including Fatty Acid to Acetyl-CoA and Acetyl-CoA to Fatty Acid, which was consistent with the absorption and utilization of volatile fatty acids in rumen fermentation (Extended Data Fig. 9f). In contrast, the reticulum epithelium showed lower metabolic fluxes compared to other chambers, suggesting a limited role in downstream energy conversion and overall lower metabolic activity.

### Differentiation trajectory of epithelial cells in forestomach

Pseudotime analysis of forestomach epithelial cells revealed a unidirectional differentiation trajectory, from pro_basal to basal, spinous, and finally granular cell subtypes (Fig. 4g and Extended Data Fig. 9c). The genes with dynamic expressions are grouped into three pseudo-chronological clusters C1–C3 (Fig. 4h). The cluster C1 is characterized by high expression early in the epithelial differentiation trajectory, and the 266 genes in C1 are significantly enriched in functions related to the regulation of cell proliferation (e.g., *MKI67*^63^, *TOP2A*^64^; Supplementary Table 9), as evidenced by the enriched GO terms of nuclear division and kinetochore assembly (Fig. 4h). The 1169 genes in C2 with high expression in the mid-stage epithelial cell development are enriched in cell growth and morphogenesis regulation (Fig. 4h). C3 (161 genes) is active in the late-stage of epithelial cell development, and contains genes associated with epithelial keratinization (e.g., *IVL*^65^, *KRT6A*; Supplementary Table 9), as the GO terms include keratinization and keratinocyte differentiation (Fig. 4h). The expression of *KRT6A*, a marker for keratinization^66^, presented a significant response to the differentiation trajectory of epithelial cell keratinization (Fig. 4h).

### Divergent evolutionary origins of forestomach and abomasum

We performed evolutionary origin analysis to investigate the evolutionary basis underlying the divergence between the forestomach and abomasum. Firstly, we conducted a comparative analysis of publicly available intestine scRNA-seq data from zebrafish^67^ with our scRNA-seq data in glandular stomach of turkey (two-chambered avian species), horse (monogastric mammal) stomach, and abomasum of camel (three-chambered ruminant) and buffalo (four-chambered ruminant). It demonstrated remarkable conservation of cell types across these species, suggesting fundamental evolutionary constraints on digestive system development (Fig. 4i). The observed similarity cell types in the zebrafish intestine and the stomachs of these species supported the fact that the vertebrate stomach evolved from the ancient intestine expansion^68^. However, a large portion of cells cannot be assigned with any cell type in zebrafish intestines due to a lack of cell-type marker genes, suggesting that the gene expression profiles of these vertebrates’ stomachs have undergone extensive remodeling since their divergence from the zebrafish lineage. A pronounced cross-species mutual conversion of epithelial and B cells indicates their relatively evolutionary instability in vertebrates.

Additionally, we also compared the scRNA-seq data of human esophagus^69^ with the rumen and abomasum in ruminants. We revealed that enteroendocrine cells in the abomasum predominantly originate from esophageal fibroblast, and other cell types such as monocytes and B cells also exhibit notably distinct lineage patterns (Fig. 4j). Also, the human esophagus shared more expressed genes with rumen (deer: *n* = 383; buffalo: *n* = 912), comparted to abomasum (deer: *n* = 79; buffalo: *n* = 2) (Fig. 4k). Collectively, our results supported that the forestomach originated from the esophagus^4^ and the abomasum evolved from intestinal structures^68^.

### Evolutionary adaptations of vertebrate stomachs to diet plant diversification

The evolutionary trajectory of vertebrate stomach is intricately linked to major ecological transitions and the diversification of plant species, which are the primary dietary source for vertebrates (Fig. 5). The forces by plant evolution, ecological change, and digestive innovation have driven the formation and diversity of stomach in aves (e.g., two stomach chambers), monogastric mammals (e.g., three feeding habits), and ruminants (e.g., 3–4 stomach chambers). The gastric glands first appeared approximately 350–450 Mya, which coincided with the origin of vascular plants^70^. The bird-mammal split around 318 Mya^71^, laying the foundation for divergent digestive adaptations which corresponded to the boom of ferns and the emergence of gymnosperms. The digestive systems in birds and mammals underwent distinct evolutionary paths, and diversified shortly after the Cretaceous–Paleogene (K-Pg) boundary around 66 Mya, following the mass extinction of dinosaurs^72–75^. Sharing the same ancestor to the birds, some herbivorous dinosaurs were believed to use gastroliths for mechanical digestion^76^, much like the function to the muscular stomach of modern birds. With the emergence of avian muscular stomach, muscular stomach-specific expression of *COL6A3*, which strengthens extracellular microfibril formation^77^, could have supported its evolution for mechanically digesting hard angiosperm seeds and fruits (Fig. 5). The global shift toward cooler, drier climates around 40 Mya coincided with the emergence of a fully functional rumen^78^. This evolutionary event likely served as an adaptation to the concurrent expansion of coarse and fibrous grasses (Poaceae), establishing the digestive foundation for their utilization. Following the emergence of rumen, rumen-specific expression of *JCHAIN*, which is crucial for establishing the immune balance necessary for maintaining the complex microbial fermentation in rumen^79^, likely facilitated the digestion of coarse grasses for ruminants (Fig. 5). Processing coarse fibrous plants necessitates robust gastric motility, which is fundamentally driven by the contractile activity of SMCs^80^. *LUC7L* is specifically expressed in SMCs and plays a critical role in maintaining the contractile property (Fig. 5). These adaptations are essential for the rumination that enables the efficient propulsion and mixing of fibrous digesta across the multiple chambers.

**Fig. 5.**
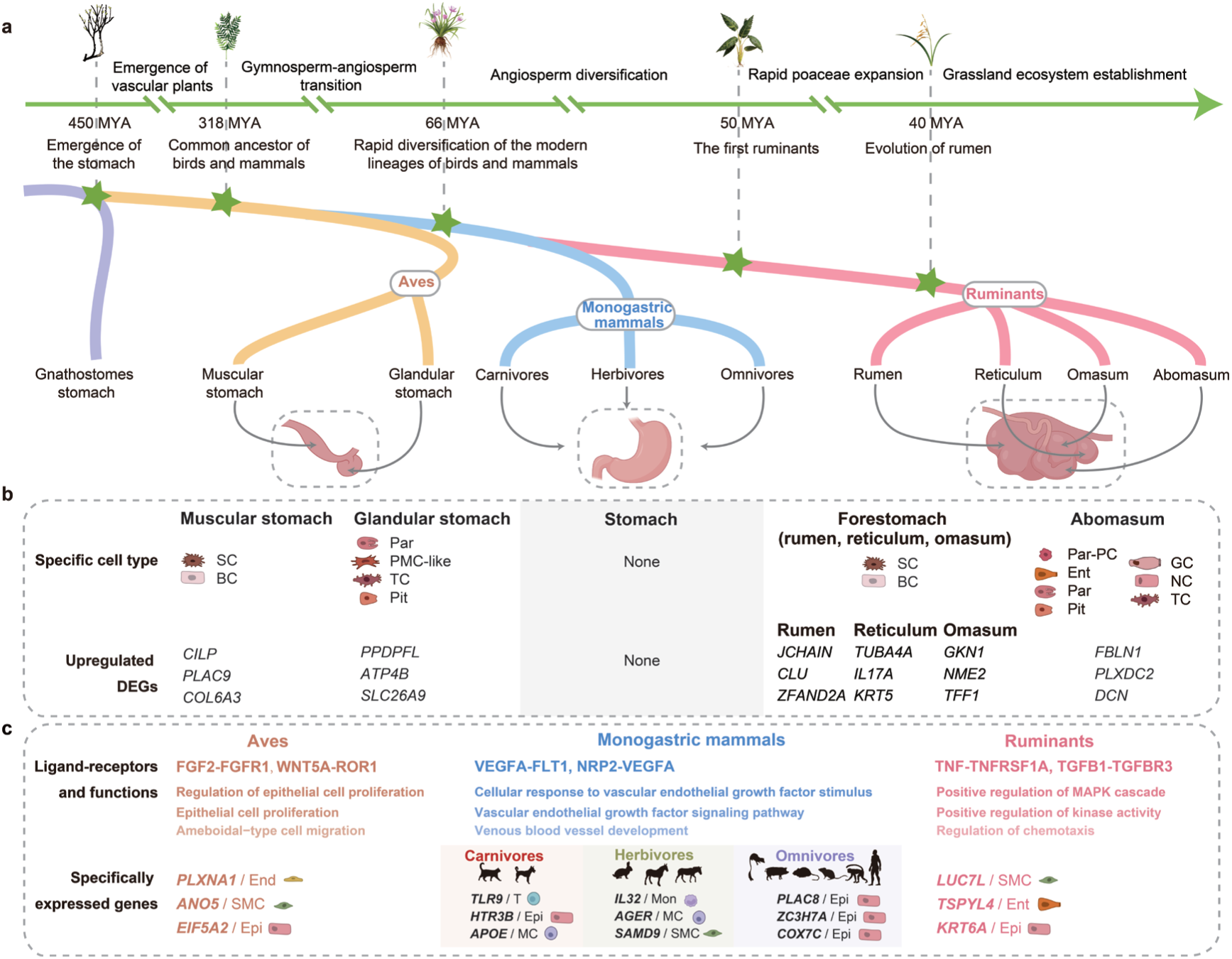
Overview of vertebrate stomach evolution along the diversification of diet plant species. a, Evolutionary timeline of the vertebrate stomach. Key events trace a pathway from the emergence of the gnathostomes stomach to the stomach diversification of modern avian (i.e., muscular and glandular stomachs) and mammalian (e.g., rumen, reticulum, omasum, and abomasum of ruminants) lineages. The stomach evolutionary events are tightly coupled with the evolution of land plant. MYA, million years ago. **b,** Divergence of cell types and genes across the stomach chambers in polygastric animals. Unique cell types and the differentially expressed genes (DEGs) for a given stomach chamber were determined by comparison with all other chambers of the same species. **c,** Divergence of cell types, ligand–receptor pairs, and genes in aves, monogastric mammals, and ruminants. The full names of cell types are shown in Supplementary Table 17.

### Spatial heterogeneity of cell types and gene expressions for herbivore stomach

To characterize the spatial patterns of cell types and gene expression, we performed single-cell spatial transcriptome analysis (Stereo-seq) of 10 longitudinal sections covering the four stomach chambers of two sheep and the stomachs of two horses (Fig. 1a). Based on the identified cell types in our scRNA-seq data, we annotated cell types in the spatial map of Stereo-seq. We applied a recently developed method, “Cell2location”, to obtain the bin50 spots on the Stereo-seq map (Fig. 6a). AUROC correlation analysis demonstrated a high correlation between gene expression patterns identified by Stereo-seq and those by scRNA-seq (Extended Data Fig. 10b). The cell-type distribution was consistent between the replicates of both sheep and horses (Fig. 6a and Extended Data Fig. 10a).

**Fig. 6.**
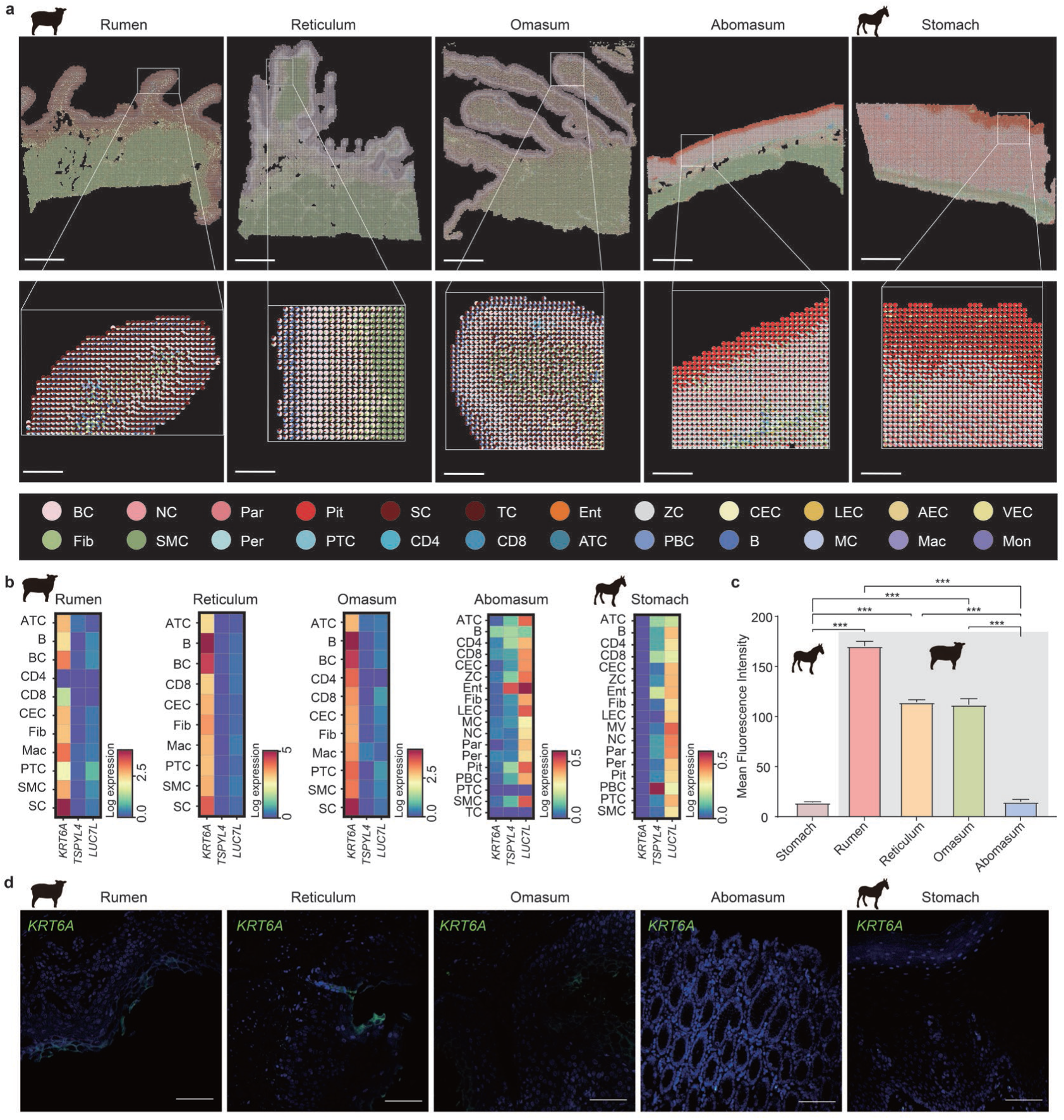
Spatial transcriptomic patterns among ruminant stomach chambers. a, Stereo-seq maps (top) of a stomach section across the four stomach chambers (rumen, reticulum, omasum, and abomasum) of sheep and the equine stomach, showing the spatial distribution of different cell types, with a scale bar of 1 mm. The boxed region of each section is shown at a higher resolution (bottom), with a scale bar of 200 μm. **b,** Heatmap of the mean log-normalized expression of *KRT6A*, *TSPYL4*, *LUC7L* in each cell type across four stomach chambers of sheep and equine stomach. **c,** Statistical analysis of the average fluorescence intensity of *KRT6A* are performed in four stomach chambers of sheep and equine stomach. Data are presented as the mean ± SEM at three independent experiments. *P* values are determined by Wilcoxon test, where *** indicates *P* < 0.001. **d,** Immunofluorescence staining of *KRT6A* in four stomach chambers of sheep and equine stomach, with a scale bar of 100 μm. The full names of cell types are shown in Supplementary Table 17.

The resultant spatial maps revealed fine-grained anatomical structures of the stomachs and localized major cell types including epithelial, endothelial, stromal and immune cells in the four chambers of sheep and the stomachs of horses (Fig. 6a and Extended Data Fig. 10a). Specifically, we observed distinct papillary structures in forestomach that significantly increase the surface area to enhance absorption and mechanical friction^81^. These regions are predominantly composed of basal and spinous cells, which contribute to the protective and abrasive functions of forestomach. Also, basal cells were spatially distributed close to the spinous cells (Fig. 6a), with their peaks at the outermost layer towards the stomach contents in forestomach (Extended Data Fig. 10c). In contrast, the abomasum exhibited well-defined gastric folds and pits, which are lined with pit cells and various secretory cells such as parietal cells, enteroendocrine cell, and tuft cells, supporting its role in enzymatic digestion and acid secretion (Fig. 6a and Extended Data Fig. 10a). In abomasum and the equine stomach, pit cells are closest to the stomach contents, and the peaks of neck and chief cells were located at the inner epithelial layer (Extended Data Fig. 10c). Immune cells showed a closely intermingled distribution with epithelial cells within the papillary structures of forestomach, which was consistent with their enrichment in forestomach based on scRNA-seq (Figs. 4b, 5a and Extended Data Fig. 10a). We observed pronounced colocalization between spinous and T cells in forestomach, and between tuft and pit cells in abomasum and equine stomach (Extended Data Fig. 10d). Thus, the spatial organization of various cell types is highly structured in the four chambers of sheep and equine stomach, reflecting their adaptation to different digestive and absorptive functions.

We next investigated the spatial expression of *KRT6A*, *TSPYL4*, and *LUC7L* across all the cell types. We observed a high expression of *KRT6A* in spinous cells of forestomach compared to abomasum and equine stomach (Fig. 6b and Extended Data Fig. 11a), as also indicated by the immunofluorescence results (Fig. 6c, d). *KRT6A* expression decrease from the stomach content side to the outer surface followed the distribution of the spinous cells (Extended Data Fig 11c). A high expression of *LUC7L* was identified in muscular layers in the four stomach chambers (Extended Data Fig. 11a), particularly in SMCs, whereas this pattern was not observed in equine stomach (Fig. 6b). Among the four stomach chambers, the highest expression of *LUC7L* was found in abomasum, as confirmed by fluorescence in situ hybridization results (FISH) (Fig. 7a). *TSPYL4* expression was the highest in abomasum, which agreed with the spatial expression results (Fig. 6b and Extended Data Fig. 11b). Taken together, these genes might be involved in regulating key functional activities in the four stomach chambers of ruminants.

**Fig. 7.**
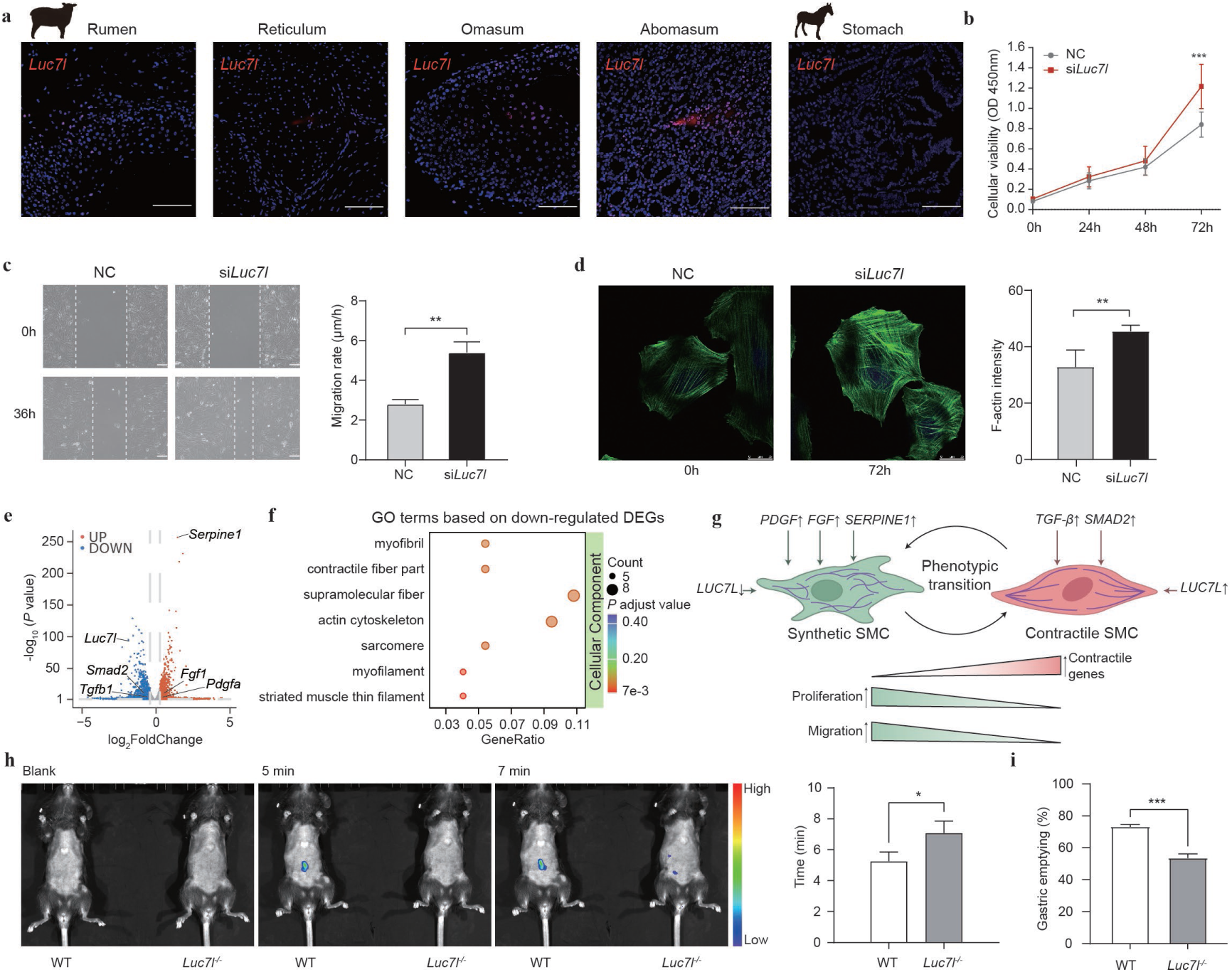
Impact of *Luc7l* knockdown on A7r5 cells and gastric motility in mice. a, In situ hybridization of *LUC7L* transcripts in four stomach chambers (rumen, reticulum, omasum, and abomasum) of sheep and equine stomach, with a scale bar of 100 μm. **b,** Quantification of the cell proliferation between negative control (NC) and *Luc7l*-knockdown (si*Luc7l*) A7r5 cells, as measured by CCK-8 assay. Data are presented as the mean ± SEM (3 biological replicates). *P* values are determined by two-way ANOVA, where *** indicates *P* < 0.001. **c,** Migration of A7r5 cells in a comparison of NC and si*Luc7l*. Wound healing of A7r5 cells was assessed in images (left) at 36 hours (h) post-scratch, relative to the 0-h time point. Wound edge is highlighted by the dashed white line. Migration rate was calculated and compared between NC and si*Luc7l*. Data are presented as the mean ± SEM (3 biological replicates). *P* values are determined by two-tailed unpaired *t*-test, where ** indicates *P* < 0.01, with a scale bar of 100 μm. **d,** Cytoskeleton F-actin of A7r5 cells in a comparison of NC (at 0 h) and si*Luc7l* (at 72 h). The images were all taken from randomly selected fields. Data are presented as the mean ± SEM (3 biological replicates). *P* values are determined by two-tailed unpaired *t*-test, where ** indicates *P* < 0.01, with a scale bar of 25 µm. **e,** Volcano plot showing the up- and down-regulated differentially expressed genes (DEGs) in bulk RNA-seq between the NC and si*Luc7l*. **f,** GO cellular component (CC) enrichment analysis of the down-regulated DEGs. **g,** Schematic diagram illustrating the mechanism that *LUC7L* regulates the phenotypes of smooth muscle cells (SMCs). The up-regulated expression of *LUC7L* promotes the switch from synthetic to contractile phenotypes and lowers rates of SMC proliferation and migration. The expression of synthetic genes (e.g., *PDGF*, *FGF*, and *SERPINE1* genes) are up-regulated significantly in si*Luc7l* cells, while the contractile genes (e.g., *TGF-β* and *SMAD2*) are down regulated, compared to NC. **h,** Representative ventral images of fluorescence emission of abdomen. Mice received a 600 mg/kg body weight of 4-kDa FITC dextran imaging agent in wild-type (WT) mice and homozygous *Luc7l* knockout (*Luc7l*^−/−^) mice. Representative ventral images of anesthetized mice are shown at baseline (blank), 5 and 7 min post-gavage. All images were acquired using the same pseudocolor scale of radiance to show relative changes in bioluminescence emission over time. Time to first appearance of fluorescence in the abdomen compared between WT and *Luc7l*^−/−^ mice. Data are presented as the mean ± SEM (3 biological replicates). *P* values are determined by two-tailed unpaired t-test, where * indicates *P* < 0.05. **i,** Gastric emptying 20 min after gavage of 0.5 ml of phenol red solution in WT and *Luc7l*^−/−^ mice. Data are presented as the mean ± SEM (6 biological replicates). *P* values are determined by two-tailed unpaired *t*-test, where *** indicates *P* < 0.001.

### Regulatory effects of *Luc7l* gene on smooth muscle cells and gastric motility

Previous studies have shown that mouse *Luc7l* is related to skeletal muscle growth and development^35,82^. To elucidate the function of *Luc7l*, we generated *Luc7l*-knockdown rat smooth muscle cells (A7r5) using small interfering RNA (siRNA) (Extended Data Fig. 12b). Our findings revealed that the *Luc7l* knockdown significantly promoted the proliferation and migration of A7r5 cells (Fig. 7b, c and Extended Data Fig. 12c). This observation was further corroborated by immunofluorescence analysis of F-actin (Fig. 7d and Extended Data Fig. 12d). The volcano plot delineates the transcriptomic changes following the *Luc7l* knockdown, identifying 2660 up-regulated (e.g., *Serpine1* and *Lgmn*), and 2377 down-regulated genes (e.g., *Atp5mc3*, *Arpc5*, and *Luc7l*) (Fig. 7e and Supplementary Tables 10, 11).

Further comparative transcriptomic analysis between the *Luc7l*-knockdown cells and negative control cells revealed significant enrichment of up-regulated genes in pathways related to proliferation, while down-regulated genes in pathways related to muscle contractile function (Fig. 7f and Extended Data Fig. 12e, f). These findings showed that reduced *LUC7L* expression leads to upregulation of *PDGF*^83^, *FGF*^84^, and *SERPINE1*^85^ (marker genes of the synthetic phenotype), while high *LUC7L* expression correlates with elevated expressions of *TGF-β* and *SMAD2*^84^ (marker genes of the contractile phenotype) (Supplementary Table 10). The phenotypic switch from synthetic to contractile was driven by the suppression of proliferation and migration (Fig. 7b and c) and transcriptional reprogramming of marker genes in smooth muscle cells (Fig. 7g).

To investigate the role of *Luc7l* in gastric motility, we generated homozygous *Luc7l* knockout (*Luc7l*^−/−^) mice using CRISPR-Cas9 technology and used littermate wild-type (WT) mice as controls (Extended Data Fig. 13a and b). *In vivo* fluorescence imaging was performed following intragastric administration of a FITC-labeled substrate (Fig. 7h). At 5 min after gavage, clear FITC fluorescence signals were detected in the intestinal region of WT mice, whereas no intestinal fluorescence was observed in *Luc7l*^−/−^ mice. By 7 min post-gavage, only weak intestinal fluorescence signals were detected in *Luc7l*^−/−^ mice, indicating a markedly slower transit of gastric contents to the intestine compared with the WT mice. Considering the time to first detection across all individuals, FITC fluorescence appeared in the intestine significantly later (*P* < 0.05) in *Luc7l*^−/−^ mice (7.08 ± 0.78 min) than in WT mice (5.25 ± 0.64 min), indicating a substantial delay in the initiation of gastric emptying (Fig. 7h). To further quantify this phenotypic difference, we measured the gastric emptying rate of the WT and *Luc7l*^−/−^ mice (Fig. 7i and Extended Data Fig. 13c).

*Luc7l*^−/−^ mice exhibited a significantly reduced (*P* < 0.001) gastric emptying rate (53.97% ± 6.22%) compared to WT mice (73.65% ± 3.33%), exhibiting significantly reduced gastric emptying relative to the WT controls, demonstrating impaired gastric motility. Collectively, these findings indicate that *LUC7L* plays a critical role in the coordinated multi-chambered gastric motility that underlies rumination in ruminants.

## DISCUSSION

In this study, we presented the most comprehensive cellular transcriptomic atlas of vertebrate stomachs, encompassing both monogastric and polygastric species across herbivorous, carnivorous and omnivorous feeding habits. Evolution of the stomach involved functional divergence and the emergence of new chambers. Our studies leveraged the combined power of scRNA-seq and Stereo-seq to construct a comprehensive cell atlas of vertebrate monogastric and polygastric stomachs at molecular and spatial single-cell resolution. We profiled the differences in cell-type abundances and cell-specific gene expressions among monogastric and polygastric vertebrates with differentiated digestive adaptation to various dietary patterns. In particular, we characterized the transcriptional profiles and spatial expression patterns of individual cells across herbivorous stomachs, including the four chambers of ruminants and the stomach of monogastric animals. Our findings elucidated the molecular foundations of multi-chambered stomach evolution and its adaptation to fiber-rich diets in ruminants via rumination.

The evolution of the vertebrate stomach is closely intertwined with that of terrestrial plants, driven by plant-herbivore co-evolutionary dynamics^86,87^. Our timeline of gastric evolution corroborates this, illustrating that major ecological transitions and dietary shifts continuously drove the specific molecular and cellular adaptations of the stomachs over time (Fig. 5). Diverse vertebrates, such as ruminants and birds, have evolved specific cell types and multi-chambered stomach structures to process tough plant materials, enabling highly efficient mechanical grinding and microbial fermentation^4,14,90,91^. Ruminants have evolved multi chambered stomachs, which supports high fiber herbivory through forestomach microbial fermentation, coinciding with grassland expansions around 40 Mya and providing advantages over monogastric systems for coarse vegetation^4^. In contrast, birds developed a two chambered stomach (glandular stomach for chemical digestion and muscular stomach for mechanical grinding), compensating for their toothlessness and facilitating breakdown of hard seeds, fruits, or fibrous plants as an evolutionary adaptation to vertebrate plants^91^.

Previous studies have revealed that dietary specialization has driven genomic changes across mammals. For instance, carnivore genomes exhibit shared evolutionary adaptations in genes (e.g., *GAP43* and *PTPN20*; Supplementary Table 7) associated with diet^88^, while both herbivores and carnivores demonstrate convergent losses of genes (e.g., *SYCN* and *NOX1*) whose functions are no longer essential for their respective diets^89^. Building upon these genomic insights, our cross-species atlas provided additional cellular insights into the evolution of vertebrate stomachs associated with diverse diets.

We find that each cell type possesses distinct evolutionary histories in gene expression and regulatory features in the stomachs across species. Endothelial cells displayed the highest evolutionary conservation among all cell types (Fig. 2a). This conservation is necessary for maintaining vascular integrity and tissue homeostasis^92^, and might facilitate the enhanced absorption and transport of nutrients^93^. Also, the evolutionary conservation of cell types is of great significance for gastric disease, because rapidly evolving genomic regions influence disease risk^94^ and are linked to metabolic and gastrointestinal disorders^95–99^. The slower evolutionary rate of endothelial cells in ruminant stomachs could have contributed to their lower susceptibility to gastric cancer compared to the other species. In contrast, immune cells are among the most rapidly evolving cell types across animal tissues^11,100^, which reflects their need to adapt to the unique immune challenges of the vertebrate stomach^101^. Notably, most ruminant stomach cell types have higher evolutionary rates, implying the rapid evolution of their multi-chambered stomachs.

We show that the profound differences in cellular composition and gene expression between the ruminant forestomach and abomasum, beyond their morphological distinction, stem from their distinct origins^4,59,68^. The forestomachs are specialized for microbial fermentation and mechanical processing of fibrous forage^102^. Particularly, the rumen epithelium, characterized by a more complex cell type composition and a higher relative abundance, may potentially drive changes in the microbial ecosystem and influence metabolic pathways associated with methanogenesis^103^. Although both horses and sheep primarily feed on herbaceous plants, their differing forage preferences, including horses favoring high-quality grasses (approximately 70–90%) and sheep consuming a more even proportion of grasses (approximately 50–60%), forbs, and legumes^104,105^, may contribute to differences in stomach structure and cellular composition.

Of note, *KRT6A*, a type II keratin protein, is a critical component of cytoskeleton in mammalian cells^106^, and highly expressed in the spinous cells of the forestomach. As keratins provide mechanical strength and resilience and enable epithelial cells to withstand physical stress and prevent damage^107^, *KRT6A* is involved in epidermalization of squamous epithelium and plays an important role in cell growth, proliferation and migration, especially keratinocyte migration^108,109^, in the ruminant forestomach. The abomasum, which resembles the monogastric stomach^59,110^, contains unique epithelial cell types (e.g., enteroendocrine and pit cells) that are absent from the forestomach and likely support its secretory functions^111^. Also, *TSPYL4* is highly and specifically expressed in the enteroendocrine cells of abomasum. The TSPYL family genes (*TSPYL1*-*TSPYL6*), which are involved in chromatin remodeling and transcriptional regulation^112–115^, may fine-tune the chemical digestive processes of the abomasum. We also observed the higher expression of *LUC7L*, a serine/arginine-rich protein, in ruminant stomach smooth muscle compared to monogastric herbivores. The teeth-less aves have evolved a muscular stomach to mechanically break food, which represents enhanced expression of *LUC7L* at the single-cell level (Extended Data Fig. 12a), suggesting a molecular reinforcement of its function in muscle contraction. As this gene is downregulated transcriptionally during muscle differentiation^35^, its knockdown triggered a phenotypic shift from a mature, contractile state to a less differentiated, proliferative one. Given that gastrointestinal motility relies on coordinated smooth muscle contraction^116^, *LUC7L* likely helps maintain the contractile phenotype to stabilize rhythmic contractions and prevent dysmotility, which may be essential for multi-chambered morphology of ruminants.

Our findings not only reveal the cellular and molecular basis of gastric evolutionary adaptation but also provide insights for enhancing cellulose conversion and utilization efficiency through animal genetic engineering. For example, the key genes and regulatory networks identified in this study in the ruminant stomach—including the *KRT6A* gene that enhances mechanical resistance in the forestomach epithelium, the *LUC7L* gene essential for maintaining smooth muscle contractility in multi-chambered stomachs—offer potential genetic targets for engineering monogastric animals to acquire ruminant-like digestive capabilities. By introducing or enhancing the expression of these ruminant-specific genetic elements, it may be possible to transform the monogastric structure of non-ruminants (e.g., swine and horse) into a multi-chambered system functionally resembling the “rumen-reticulum-omasum-abomasum”. Such modifications could enable monogastric animals to efficiently break and utilize fibrous plant materials, significantly improving their cellulose utilization efficiency while reducing dependence on high-quality feed. This approach holds important practical implications for sustainable livestock production and efficient resource utilization. Also, the gene *LUC7L*, a crucial regulator of gastric motility, represents a potential therapeutic target for its associated disorders.

In conclusion, our study uses a cross-species scRNA-seq approach coupled with Stereo-seq to comprehensively investigate the molecular evolution of stomach. Our data and results provide an excellent reference and a valuable resource for deciphering the molecular and cellular basis underlying the stomach biology across vertebrates.

### Limitations of the study

A couple of limitations are noted in this study. First, biological replicates were not included for some species. Nevertheless, the inclusion of multiple species within each of the three evolutionary animal groups (i.e., ruminants, monogastric mammals, and birds) effectively serves as biological replicates at the group level, thereby enhancing the robustness and generalizability of our cross-species comparative analyses. Also, rumination-relevant functions of the three identified genes in specific cells probably require further validation in the large monogastric animal models like swine. Future research by integrating single-cell multi-omics data with single cell imaging technologies would surely deepen our understanding of molecular mechanisms underlying the development and specification of various stomach chambers across vertebrates including human.

## METHODS

### Ethics statement

All sample collection was approved by the Ethics Committee of China Agricultural University (CAU20160628-2). All sampling procedures strictly followed the animal experimental guidelines issued by the Ministry of Science and Technology of China, and complied with the Convention on Biological Diversity and the Convention on Trade in Endangered Species of Wild Fauna and Flora.

### Sample collection for scRNA-seq

Stomach tissue samples (*n* = 38) from 21 vertebrate species, categorized by the number of stomach chambers and feeding habit (Fig. 1a), were collected for scRNA-seq. These included monogastric animals such as two rodent species (*Rattus norvegicus*, brown rat and *Mus musculus,* house mouse), two carnivora species (*Felis catus*, cat and *Canis familiaris*, dog), one model animal (*Tupaia belangeri*, northern tree shrew), and four livestock species (*Equus ferus caballus*, horse; *Equus asinus*, donkey; *Sus scrofa*, swine; and *Oryctolagus cuniculus*, European rabbit).

Two-chambered stomach vertebrates included five avian species (*Gallus gallus*, chicken; *Meleagris gallopavo*, turkey; *Coturnix japonica*, coturnix quail; *Anas platyrhynchos*, mallard duck; and *Columba livia*, rock dove). Samples were also obtained from a three-chambered stomach species like camel (*Camelus bactrianus*, bactrian camel), and five ruminant livestock species with four-chambered stomachs (*Capra hircus*, goat; *Ovis aries*, sheep; *Bos grunniens*, yak; *Bubalus bubalis*, buffalo; *Bos taurus*, cattle; and *Cervus nippon*, sika deer). These animals are categorized into omnivorous (i.e., rat, mouse, tree shrew, swine, chicken, turkey, quail, duck and dove), carnivorous (i.e., cat and dog) and herbivorous (i.e., rabbit, horse, donkey, goat, sheep, yak, buffalo, cattle, deer and camel) species in terms of feeding habit.

The camel samples were kindly provided by the Inner Mongolia Camel Research Institute, and the northern tree shrew samples were from the Kunming Institute of Zoology of the Chinese Academy of Sciences. The other samples were collected from local markets under the permission from the Ethics Committee of China Agricultural University (CAU20160628-2).

To capture functional diversity of the whole stomach (Fig. 1a), gastric tissues from distinct anatomical locations were isolated and placed on ice immediately after sacrifice by the carotid artery exsanguination. In brief, one to four locations were sampled from each stomach or stomach chamber of an animal, i.e., four locations (cardia, fundus, body, and pylorus) in the monogastric stomach, four (dorsal sac, ventral sac, caudodorsal blind sac, and caudoventral blind sac) in rumen^117^, one in reticulum, two (body and canal) in omasum^118^, and four (cardia, fundus, body, and pylorus) in abomasum^119,120^. For the aves, we collected the tissues of muscular and glandular stomach, respectively. Approximately 5-mm-sized tissue pieces were washed with phosphate-buffered saline (PBS), and then transferred into preservation solution (Miltenyi Biotec) in 2 mL cryovials (Corning, 430917). Tissues were stored at 4℃ within 24 h before library construction. Meanwhile, duplicate tissues were fixed in 4% paraformaldehyde (Solarbio, P1110) for 24 hours at 4℃ and then embedded in paraffin for the hematoxylin-eosin (HE) staining, immunofluorescence staining and fluorescence in situ hybridization (FISH) analyses.

### Sample collection for Stereo-seq

For the Stereo-seq experiments, two adult sheep (approximately 1.5 years old, female) and two horses (approximately 2.5 years old, male) were utilized (Supplementary Table 12). The animals were sacrificed by the carotid artery exsanguination. A total of eight sections were sampled from the two four-chambered sheep, and two sections were obtained from the two monogastric horses. Following collection, tissue samples were immediately snap-frozen and embedded in liquid-nitrogen-prechilled isopentane in Tissue-Tek OCT (Sakura Finetek, 4583) and subsequently stored at −80℃ until further processing. Tissue cryosections (10 μm thickness) were prepared using a Leika CM3050S cryostat, and subsequently mounted onto Stereo-seq chip (BGI) at −20℃. The mounted chips were incubated at 37℃ for 3 min, and subsequently the chips and the tissue sections were fixed in pre-cooled methanol at −20℃ for 40 min before Stereo-seq library preparation. The remaining sections were collected for nucleic acid staining to verify tissue integrity and nucleus locations prior to in situ RNA capture. Nucleic acid staining was performed using a nucleic acid dye (Thermo Fisher Scientific, Q10212), and fluorescence imaging was obtained with a Stereo Go Optical microscope (BGI).

### scRNA-seq library construction and sequencing

For scRNA-seq, the tissues were further cut into approximately 1-mm-sized pieces and incubated in RPMI-1640 medium (Thermo Fisher Scientific, 11875500137) at 37℃ for 30 min. Following centrifugation at 300*g* for 5 min, the precipitates were digested in RPMI-1640 medium mixing with 2.5 mg/mL collagenaseⅡ (Solarbio, C8150) and 2 kU/mL DNase I (Solarbio, D8070) at 37℃ for 30–60 min to yield data of adequate quality (Supplementary Table 1). The resulting cell suspension was filtered through a 70-µm sterile cell strainer (Thermo Fisher Scientific, 22363548) to remove large tissue debris and cell aggregates. After centrifugation at 300*g* for 5 min at 4℃, the supernatant was discarded, and the cell precipitates were washed twice with PBS by vortexing and centrifugation. Afterwards, an automated cell counter (Luna-II, Logos Biosystems) was used to estimate the number of live cells and the Rigel S3 fluorescence cell analyzer (Countstar) was used to assess the cell viability and concentration. Cell viability > 80% was set as the threshold and cell suspension concentration was adjusted to 300–600 living cells per microliter. To enhance the efficiency of sorting robust and live cells for single-cell experiments, the MACS® Dead Cell Removal Kit (Miltenyi Biotec) was used to eliminate the dead cells. To achieve the concentration needed for scRNA-seq, the sorted single-cell suspension was counted and diluted to the final concentration in DMEM (Thermo Fisher Scientific, 11320-033) with different volumes (500–1,000 µL) of 10% fetal bovine serum (FBS) (Solarbio, s9020). Cell suspensions from distinct stomach locations were thoroughly mixed in equal proportions and used for scRNA-seq library construction. For comparative analysis of multi-chambered stomachs, we constructed scRNA-seq libraries for separate cell suspensions from: (1) muscular and glandular stomach of turkey; (2) rumen, reticulum, and abomasum of camel; and (3) rumen, reticulum, omasum, and abomasum of deer, buffalo, and sheep (Fig. 1a).

The cell suspensions containing at least 8000 cells were loaded on a Chromium Single Cell Controller (10X Genomics) to generate single-cell gel bead-in-emulsion (GEM), using Single Cell 3’ Library and Gel Bead Kit V3 (10X Genomics, PN1000350) and Chromium Single Cell Chip Kit (10X Genomics, PN120236) according to the manufacturer’s protocol. Briefly, thousands of suspended cells were captured into GEM containing 10X cell barcodes, unique molecular identifier (UMI), and poly-dT primers. The released RNA from lysed cells was used for barcoded full-length complementary DNA (cDNA) amplification in a C1000 Touch Thermal Cycler (Bio-Rad) with 12 PCR cycles of 53℃ for 45 min and 85℃ for 5 min. The cDNA was cleaned using Dynabeads MyOne Silane beads (Thermo Fisher Scientific, 37002D), and was further amplified on a Biometra TProfessional Thermocycler (Montreal Biotech). Sequencing was performed on a Novaseq6000 (Illumina) with a depth of > 100,000 reads per cell and pair-end 150 bp (PE150).

### Stereo-seq library construction and sequencing

For capturing RNAs, tissue slices for Stereo-seq chips were treated using 0.1% pepsin (Sigma-Aldrich) in 0.01 M HCl buffer and incubated at 37℃ for 5 min. RNA released from the permeabilized tissues and captured by the DNA nanoballs (DNBs) on Stereo-seq chips was reverse transcribed overnight at 42℃using reverse transcriptase SuperScript II (Invitrogen, 18064-014). After reverse transcription, Stereo-seq chips were washed twice with wash buffer and digested with tissue removal buffer (10 mM Tris-HCl, 25 mM EDTA, 100 mM NaCl, 0.5% SDS) at 55℃ for 10 min. The cDNAs were released from the chip with the treatment of exonuclease I (New England Biolabs) over night at 55℃, and was purified using the VAHTS DNA Clean Beads (Vazyme, N411-03). The cDNA’s amplification was performed using KAPA HiFi Hotstart Ready Mix (Roche Diagnostics) following a program of initial incubation at 95℃ for 5 min, 15 cycles at 98℃ for 20 s, 58℃ for 20 s, 72℃ for 3 min and a final incubation at 72℃ for 5 min. After quantification in QubitTM dsDNA Assay Kit (Thermo Fisher Scientific, Q32854), a total of 20 ng cDNAs were then fragmented with in-house Tn5 transposase at 55℃ for 10 min, and the fragmentation reactions were terminated by adding 0.02% SDS and incubation at 37℃ for 5 min. Fragmented cDNAs were then amplified with KAPA HiFi Hotstart Ready Mix, 0.3 mM Stereo-seq-Library-F primer, and 0.3 mM Stereo-seq-Library-R primer in a total volume of 100 mL. The PCR reaction was run with a program of 95℃ 5 min; 13 cycles of 98℃ 20 s, 58℃ 20 s and 72℃ 30 s; and 72℃ 5 min. PCR products were purified using the AMPure XP Beads. The libraries were then sequenced on MGI DNBSEQ-T7 sequencer according to the standard protocols at Novogene (Tianjin, China), with 35-bp read 1 and 100-bp read 2.

### Data processing and quality control of filtered cells

Besides the data created, we downloaded stomach scRNA-seq data of human (*Homo sapiens*) (GenBank accession no. GSE134355) and crab-eating macaque (*Macaca fascicularis*) (CNGB accession no. CNP0001469), and combined the data in the integrated analyses. All the chromium scRNA-seq data was processed using the CellRanger v7.0 (https://github.com/10xGenomics/cellranger) in the 10X Genomics analysis pipelines with the default parameters. Briefly, raw sequencing data was demultiplexed into FASTQ files using the *mkfastq* function of CellRanger. Then, the reads were aligned to the corresponding reference genomes (Supplementary Table 1). The gene count matrix was generated for each GEM based on cell barcodes and UMI with a Phred quality score ≥ 30, using the *count* function.

To obtain high-quality cells, we removed cells that met the following criteria for each sample: expressing less than 200 genes (nFeature < 200) and the percentage of mitochondrial genes more than 20% (percent.mt > 20), using the R packages Seurat^121^ v4.3.0. Doublets were identified and discarded by using DoubletFinder^122^ v2.0.3. Standard preprocessing steps were applied for the gene expression data normalization using *NormalizeData*, and cell clustering was performed on the normalized gene expression data using *RunPCA*, *FindNeighbors*, and *FindClusters*. For all the animals, we set the parameters with “PC = 1:30, dimension = 1:30, and resolution = 0.8”. DEGs in each cluster were identified using *FindAllMarkers* and *FindMarkers*. The cell clusters were annotated as corresponding cell types based on expressions of the typical marker genes reported previously.

### Integration of datasets based on orthologous genes

To determine the best integration method for our datasets, we utilized scIB^123^ to benchmark several widely used Python-based tools: CCA^124^, RPCA^124^, Harmony^125^. We used 10 metrics in scIB, and the total scores were calculated as a weighted mean (40/60) of batch correction and biological variance conservation. Among the tools, CCA as the top performer was used for data integration and correction for the batch effect^126^. First, we obtained the one-to-one (1:1) orthologous across all the 23 vertebrate species (Supplementary Table 13). In brief, orthologous genes for deer, dove, and buffalo were determined against the protein sequences of humans using BLAST as implemented in Orthofinder^127^ v2.5.5. For the other 20 species, orthologous genes were retrieved from Ensembl (http://dec2021.archive.ensembl.org/index.html; Dec 2021 v105) using the R package biomaRt v2.57.1 (https://github.com/Huber-group-EMBL/biomaRt). The resulting orthologous genes were concatenated and integrated based on human proteins, yielding a total of 4,498 orthologous genes across the 23 species (Supplementary Table 13). Expression matrix of orthologous genes was retained and gene normalization was performed using *NormalizeData*. Anchors were identified by *FindIntegrationAnchors*, and normalized expression data were integrated using *IntegrateData.* Finally, a total of 3,682 orthologs with detectable expression (defined as ≥ 1 UMI in at least three cells of any cell type) were obtained (Supplementary Table 13). Cell clustering was performed using *RunPCA*, *FindNeighbors*, *FindClusters* and *RunUMAP*.

### Annotation of gastric cell types

To annotate cell types across the 23 vertebrate species, we collected known marker genes from human and mouse stomachs^15,128–130^, with one to three marker genes per cell type (Fig. 1e and Supplementary Table 3). Based on UMAP coordinates and the hierarchical clustering of cell types (Fig. 1c), we annotated a total of 9 broad cell types that consist of 34 cell subtypes (Supplementary Table 2). We evaluated the reproducibility and classification robustness of the identified cell types across species using MetaNeighbour^131^ v1.9.1. The top quartile of highly variable genes was selected using *get_variable_genes*, and the area under the receiver operator characteristic curve (AUROC) scores were calculated using *MetaNeighbourUS*. The AUROC results were employed to plot heatmaps and pairwise cell type comparisons using R package ggplot2 v3.5.1 (https://github.com/tidyverse/ggplot2) (Extended Data Fig. 2d). In addition, we identified a set of novel marker genes for these cell types using *FindAllMarkers* with the parameters “only.pos = T, min.pct = 0.25, logfc.threshold = 0.25, return.thresh = 0.05” (Supplementary Table 2).

### Identification of DEGs and GO term enrichment

To compare the global expression profiles for different cell types within one species and for one single cell type across species, the corresponding gene expression matrix was retained and integrated as clusters using *IntegrateData* of Seurat. DEGs were identified among distinct clusters using *FindMarkers*. Adjusted *P*-values were calculated using the Wilcoxon rank-sum test and Bonferroni corrections. DEGs were filtered based on the thresholds of |Log_2_ (fold change)| > 0.25 and adjusted *P*-value < 0.05. GO terms were enriched using *enrichGO* of clusterProfiler^132^ v4.10.1, and the GO terms with *P* value < 0.05 were retained.

### Global patterns of gene expression across species

Pseudo-bulk RNA-seq profiles were generated for each cell type or group of animals (e.g., monogastric mammals, aves and ruminants) using *AverageExpression* of Seurat. We performed PCA based on the normalized expression matrix of 3,682 orthologues across the 23 species, using *prcomp* in the R package stats v4.3.1. To explore evolutionary relationships, we constructed gene expression trees using the neighbor-joining approach^133^, based on pairwise expression distance matrices derived from the pseudo-bulk samples. Distances between the samples were obtained with the formula of “1 − *ρ*”, where *ρ* is the Spearman’s correlation coefficient and was calculated using *cor* of the R package stats. The neighbor-joining trees were constructed using the R package ape v5.7.1 (https://github.com/emmanuelparadis/ape), and the reliability for each branch was assessed with the 1,000 bootstrap analysis generated by randomly sampling the 3,682 orthologous genes with replacement. The tree construction and the total tree length calculations were performed using published scripts^17^ available at https://github.com/evo-bio/Spermatogenesis.

### Evolutionary forces

To explore the evolutionary forces underlying the rapid evolution of stomach, we calculated the average dN/dS values based on the longest transcripts of a set of 4,498 protein-coding orthologous genes across the 23 species. The mean dN/dS value is estimated for each cell. Orthologous protein sequences were aligned using Clustalo^125^ v1.2.4 and pal2nal^134^ v14, and dN/dS ratios were calculated using the codeml software from the PAML^135^ v4.9, based on the M0 site model with the parameters “NSites = 0, model = 0”.

### Cell-cell communication analysis

We explored cell-cell communication via ligand–receptor interactions across the 23 species using CellPhoneDB^136^ v3.0. Ligand‒receptor interaction across cell types was detected through the receptor library of human-homologous genes. The average receptor and average ligand expression were calculated as interaction score and compared among different cell types to identify their cell-specific expression. Only receptors and ligands expressed in at least 10% of cells of a given cell type were retained for further analyses, whereas an interaction was considered nonexistent if either the ligand or the receptor was unqualified. The number of ligand‒receptor pairs for each species was visualized using Cytoscape^137^ v3.7.1.

### Gene regulatory network of transcription factors

Regulatory TFs for the 23 species were downloaded from the AnimalTFDB v3.0 database (http://bioinfo.life.hust.edu.cn/AnimalTFDB/#!/). Gene regulatory networks were generated using GENIE3^138^ v1.6.0 based on the gene expression matrices of each species and the DEGs among cell types. TFs and their target genes (regulons) were identified, and enriched TF-binding motifs were predicted using genome-wide rankings based on the hg38 RcisTarget database (https://resources.aertslab.org/cistarget/), following the protocols of SCENIC^139^ v1.1.2.2. The regulons related to the most conserved TFs (ATF3, GABPA, NFKB1 and STAT1) were visualized using Cytoscape^137^ v3.7.1. For each species, regulon modules were further identified based on the connection specificity index (CSI)^132^ using R package scFunctions v0.0.0.9000 (https://github.com/FloWuenne/scFunctions). Specifically, CSI values were calculated for each pair of regulons, and hierarchical clustering was used to group regulons into modules. Regulon modules were visualized using R package Pheatmap v1.0.12 (https://github.com/raivokolde/pheatmap).

### Pseudotime trajectory analysis

Pseudotime analysis was performed using the R package Slingshot^140^ v2.9.0. The global lineage structure of the assigned cell types was identified in a minimum spanning tree with *getLineages*, with manually setting up the starting and ending clusters. Smooth trajectory curves were generated using *getCurves*. The resulting pseudotime trajectories were visualized using *FeatureDimPlot* of the R package SCP v0.4.7 (https://github.com/zhanghao-njmu/SCP).

### Single-cell disease-relevance score (scDRS) calculation

We determined the enrichment of scRNA-seq cell types for human stomach disease genome-wide association studies (GWAS) traits using scDRS^141^ v.1.0.2 with default settings and the human stomach scRNA-seq data. In brief, human GWAS results were downloaded from the GWAS Catalog database (https://www.ebi.ac.uk/gwas/home). We then mapped SNPs to the human genes (GRCh37 genome) using MAGMA^142^ v.1.1.0 with default settings. The MAGMA z score of each gene was calculated based on the resulting annotations and GWAS summary statistic. Afterwards, we selected the top 1,000 genes based on their MAGMA z scores as scDRS input data, and calculated the correlations of top 1,000 genes with the human gastric cancer via scDRS. The scDRS gastric cancer score at the cell-level was calculated via scDRS in human. Also, the Seurat disease score at the cell-type level was calculated using *Addmodulescore*^143^ in Seurat based on the top 1,000 disease genes in human (scDRS predicted). The heatmap of disease scores for each cell type were plotted in all the 23 species using the R package ggplot2.

### Identification of homogenoue cell types

To correlate gene expression patterns and determine cell type homology, we performed two independent SAMap analyses using SAMap^144^ v1.0.15: one for comparing stomach (i.e., stomach in horse, glandular stomach in turkey, and abomasum in camel and buffalo) with zebrafish intestine (GenBank accession no. GSM7184372) and the other for comparing human esophagus (GenBank accession no. GSE160269) with rumen and abomasum in deer and buffalo. We first constructed a gene-gene bipartite graph between the paired species by running ‘map_genes.sh’ module of SAMap based on BLASTP strategy. The above scRNA-seq datasets were then processed and integrated using SAMap, after which mapping scores between cell types were calculated using *get_mapping_scores* and visualized using R package networkD3 v0.4 (https://christophergandrud.github.io/networkD3/).

### Construction of a single-cell metabolic flux profile

To infer the metabolic status of cells, we further analyzed the scRNA-seq data of rumen and abomasum in camel, deer, sheep and water buffalo, the graph neural network model-based single-cell flux estimation analysis (scFEA)^62^ was used to identify the single-cell metabolic flux profiles. A total of 168 metabolic modules with 22 supermodule were downloaded from the algorithm’s official GitHub page (https://github.com/changwn/scFEA). The default parameters were utilized.

### Stereo-seq analysis

#### Stereo-seq data processing

Processing of the raw sequencing data was performed following the Stereo-seq Analysis Workflow (SAW v8.1.0, https://github.com/BGIResearch/SAW). Coordinate ID (CID for spatial positions on the chips, 1–25 bp) and molecular identifier (MID for mRNA labeling, 26–35 bp) are contained in the read 1, and the cDNA sequences are in the read 2. The CID sequences were first mapped to the predefined coordinates on the in situ captured chip, allowing a tolerance of one base pair mismatch. The MID sequences are filtered to exclude the reads containing either ambiguous bases (N) or more than two bases with a quality score of ≤ 10. The high-quality cDNA sequences with CID and MID in their sequence readers were then aligned to the genomes of sheep (GCF_016772045.1) and horse (GCF_002863925.1) using STAR (bcSTAR v2.7.1a, https://github.com/alexdobin/STAR) and mapped reads with MAPQ > 10 were counted for their corresponding genes. UMI sequences with the same CID and gene locus were collapsed, and their alignment allows 1 mismatch. Finally, CID-containing expression matrix was generated based on gene annotations of the genomes of sheep and horse.

The UMIs from 2,500 neighboring DNBs (50 × 50 DNBs) were to create a bin50 expression matrix using Stereopy^145^ v1.6.1. Following spatial aggregation, stringent quality control was applied to ensure data reliability: pseudospots with detectable expression of fewer than 3 genes and exhibiting > 20% mitochondrial gene content were removed. The resulting high-quality expression matrix was subsequently utilized for downstream analyses. The bin50 UMI count matrix was first log-normalized and the expressions of target genes *KRT6A*, *TSPYL4* and *LUC7L* were spatially highlighted in colors (Extended Data Fig. 11a) using the command sc.pl.spatial of Scanpy^146^ v1.9.6.

#### Spatial mapping of cell types

To identify cell types in each spatial pseudospot, we conducted deconvolution analysis using the Cell2location^147^ v0.1.4 based on scRNA-seq data from the four stomach chambers of sheep and the stomach of horse. Reference cell type signatures in stomach scRNA-seq were used to build a Bayesian model and deconvolute cell types in the spatial transcriptomic data. The absolute cell abundance of individual cell types across all spatial pseudospots were estimated with the parameter of N = 20 as the expected cell abundance per spot and α_y_=20 to regularize within-experiment variation in RNA detection sensitivity. The cell types and their proportions were plotted spatially (Fig. 6a) using the command cell2location.plt.plot_spatial of Cell2location and Python package matplotlib.pyplot (https://github.com/matplotlib/matplotlib).

#### Correlation of cell types between scRNA-seq and Stereo-seq method

We assessed the reliability of cell-type annotations based on the Stereo-seq data using pyMN^131^. Briefly, we calculated a similarity matrix among cell types with *MetaNeighborUS*. Values in the matrix represent the AUROC scores among cell types in different datasets, which indicate the correlations of cell types between scRNA-seq and Stereo-seq.

#### Spatial tendency analysis

To dissect spatial expression tendency, we first defined the spinous cell layer in the ovine forestomach and the pit cell layer in the ovine abomasum and equine stomach as the regions of interest (ROI), with *Spatial domain* of SOAPy (https://github.com/LiHongCSBLab/SOAPy). We then generated a binary mask file of ROI using *get*_*mask*_ *from*_*domain*. We further calculated the shortest distance from each cell/spot to the ROI boundary (contour) using *Spatial tendency* with parameters “method = poly, radius = 1,000, location = out”. Finally, we visualized the cell composition and gene expression feature with spatial tendency using *show*_*tendency*.

#### Spatial map of cell dependencies

The spatial dependencies of cell types were estimated in the three different neighborhood area sizes^148^, i.e., colocalization within a pseudospot, immediate (radius of 1 pseudospot) and extended (radius of 15 pseudospots) neighborhood. Cell-type compositions calculated by cell2location were modelled to predict the abundances of other cell types using the MISTy framework implemented in the LIANA+^149^ v0.1.9. The importance of cell-type abundances within the immediate neighborhood (radius of 5 pseudospots) was inferred based on a Random Forest model between the predictor and the target cell types, and are interpreted as cell-type spatial dependencies.

#### Hematoxylin-eosin (HE) staining

Hematoxylin-eosin (HE) staining was implemented following the routine procedures^150^. Briefly, the paraffin-embedded tissues were cut into 5 μm sections using a rotary Leica RM2255 microtome (Leica Biosystems). The slides deparaffinized in xylene for 30 min were rehydrated in gradient alcohols, stained with hematoxylin solution for 5 min, and rinsed twice with distilled water for 5 min. To remove excess stain, the stained tissues were rinsed with 1% acid alcohol for 5 s, followed immediately by washing with 50℃ water for 5 min. The slides were stained with 1% eosin ethanol solution and dehydrated in gradient ethanol and xylene. Finally, the slides were mounted with neutral resin as the mounting medium, and brightfield photographs were taken using an optical microscope (McAudi).

#### Immunofluorescence staining

For immunofluorescence staining, paraffin-embedded sections were processed with xylene and rehydrated by gradient ethanol solutions (100%, 90%, 80%, 70%, and 50%) for 5 min each. The sections in 1× citrate-based antigen retrieval solution (Sangon Biotech) were heated in a microwave oven, and stood still for 5 min, before being at 100℃ for 5 min. After antigen retrieval, tissue sections were blocked with the Immunostaining Blocking Dilution Buffer (Sangon Biotech) for 1 h at room temperature. Subsequently, samples were incubated with primary antibody (1:500, Abcam, ab93279) overnight at 4℃. The goat anti-rabbit secondary antibody (1:1,000, Abcam, ab150077) incubation was conducted for 1 h at room temperature. The excess antibodies were removed by washing, and the nuclei were stained with DAPI (Solarbio, C0065). Fluorescent images were captured using imager microscope (Zeiss).

#### RNA in situ hybridization

Fresh stomach tissues were washed and fixed with 4% paraformaldehyde (Solarbio) at 4℃ for more than 24 h, dehydrated and embedded in paraffin. Paraffin-embedded sections mounted on the slides were deparaffinized in xylene, rehydrated through graded ethanol and processed for antigen retrieval. Endogenous peroxidase activity was blocked by incubating the slides with 3% H_2_O_2_ in the dark for 15 min, followed by PBS washes 3 times for 5 min each on shaker. The slides with pre-hybridization solution (Sigma-Aldrich) were incubated at 37℃ for 1 h. After removing the solution, hybridization solution containing 2 μM Cy5-labelled *LUC7L* probes (the reference sequence accession no. XM_027962022.3, target region: 678–707 for sheep; XM_023616767.2, target region: 638–669 for horse) and Cy3-labelled *TSPYL4* probe (XM_012182543.5, target region: 1206–1238) was applied for incubation in a hybridization oven at 42℃ overnight. After hybridization, the nuclei were stained with DAPI (Solarbio, C0065), and images were captured using the microscope (Zeiss).

#### Cell culture, siRNA transfection and qRT-PCR

A7r5 vascular smooth muscle cells (hereafter A7r5, Procell Life Science & Technology) and STC-1 mouse intestinal enteroendocrine cells (hereafter STC-1, Shanghai Jinyuan Biotechnology Co., Ltd.) were maintained in DMEM (Gibco, 11995073) supplemented with 10% fetal bovine serum (FBS) (Gibco, A5670701) and 1% penicillin-streptomycin (Beyotime, C0222). Rumen epithelial cells (Shanghai Jinyuan Biotechnology Co., Ltd., China) from Hu sheep were cultured in a specialized, proprietary medium (Shanghai Jinyuan Biotechnology Co., Ltd., China). The cells were incubated at 37℃ in a humidified atmosphere with 5% CO_2_ under standard conditions. To suppress expression of *Luc7l*, *KRT6A* and *Tspyl4*, siRNAs were synthesized by GenPharma (Supplementary Table 14). The cells were seeded in 6-well plates for 24 h, and used to transiently transfect siRNA with Transfect-Mate (GenePharma, G04009) according to the manufacturer’s protocols. For siRNA transfection, 100 nM siRNAs mixed with 5 µL Transfect-Mate were incubated at room temperature for 20 min, and transferred to each cell for 6 h. The medium was replaced with the DMEM, and the transfected cells were harvested after 48 h. Total RNA was extracted from the cells using TRIzol reagent (Invitrogen), and reverse transcription was performed using PrimeScript™ RT Reagent Kit (Takara). With the *ACTB* as internal control, qRT-PCR was performed using SYBR Green Real-Time PCR Kit (Takara), and relative gene expression level was calculated using the 2^-ΔΔCT^ method. The primer sequences used in this study were included in Supplementary Table 15.

#### Bulk RNA-seq library construction and sequencing

Total RNA was extracted from isolated cells with RNA TRIzol (Invitrogen). RNA quantity and quality were assessed using agarose gel electrophoresis and a NanoPhotometer® spectrophotometer (IMPLEN). Only high-quality RNA samples (RIN > 7.0) were used for RNA-seq. The cDNA libraries were constructed from purified mRNA following the standard Illumina TruSeq Stranded mRNA library preparation protocol. The final library quality was assessed on an Agilent Bioanalyzer 2100 system. RNA-seq was implemented on the Illumina NovaSeq platform, generating 150 bp paired-end reads.

#### Bulk RNA-seq data processing and analysis

Raw reads of fastq format were firstly processed with fastp software^151^, to remove reads containing adapter, ploy-N and low-quality reads. Paired-end clean reads were aligned to the rat reference genome mRatBN7.2 (GCF_015227675.2), the mouse reference genome GRCm38.84 (GCF_000001635.24), and the sheep reference genome ARS-UI_Ramb_v2.0 (GCF_016772045.1) using HISAT2^152^ v2.0.5. Properly paired and uniquely mapped reads were extracted using SAMtools^153^ v1.11. Gene counts were generated by the featureCounts^154^ v1.5.0 program of the Subread package. We also normalized the raw counts of genes using the fragments per kilobase of exon model per million mapped fragments (FPKM) method with an in-house script. DEGs were filtered based on the thresholds of |Log_2_ (fold change)| > 1 and adjusted *P* value < 0.05. The KEGG pathways and GO terms of DEGs were obtained using the same method as above.

#### Proteomic analysis

Cell supernatants were collected, filtered, and stored at −80℃ until proteomic analysis using Astral data-independent acquisition (DIA) as previously described^155^. Briefly, each sample in a 1.5 mL centrifuge tube was lysed with lysis buffer (6 M Urea, 100 mM TEAB, pH = 8.5), and then sonicated on ice for 5 min. After centrifugation at 12,000*g* for 15 min at 4℃, the supernatant was supplemented with 1 M dithiothreitol (Sigma-Aldrich, D9163) and incubated at 56℃ for 1 h. Alkylation was then performed in the dark with iodoacetamide (Sigma-Aldrich, I6125) for 1 h at room temperature. Protein concentration was determined using the Bradford Protein Assay Kit (Beyotime, P0006). Peptides were dissolved and separated using a Vanquish Neo UHPLC system (Thermo Fisher Scientific), using a 20-min gradient from 4% to 99% solvent B (80% acetonitrile, 0.1% formic acid), with solvent A containing 0.1% formic acid. After separation by the UHPLC system, peptides were ionized via an Easy-Spray (ESI) ion source, and finally analyzed in DIA mode on an Orbitrap Astral mass spectrometer (Thermo Fisher Scientific). Under the ion spray voltage of 2.0 kV, the first-stage mass spectrometry was obtained over a range of 380 to 980 m/z at a resolution of 240,000 (at 200 m/z), with automatic gain control (AGC) set to 500%. The parent ion window size was set to 2-Th with 300 windows, with normalized collision energy (NCE) of 25%. The second-stage acquisition range was 150 to 2,000 m/z at a resolution of 80,000 on the Astral analyzer.

Raw mass spectrometry data were processed based on DIA-NN software (Direct DIA)^156^ with default parameters. To ensure high data quality, the results were filtered using the following criteria: a false discovery rate (FDR) of ≤ 1% at both the precursor and protein levels, the presence of at least one unique peptide per protein, and the proteins detected in at least 50% of samples. Subsequently, differentially expressed proteins (DEPs) were obtained with the thresholds of |Log_2_ (fold change)| > 1 and *P* value < 0.05, using the DEP2^157^ v0.5.4.02. Enrichment analysis of GO terms and KEGG pathways was performed using Interproscan software^158^.

#### Cell proliferation and Cell migration assay

Cell proliferation was evaluated at 24 h, 48 h, and 72 h after transfection using the Cell Counting Kit-8 (CCK-8, Dojindo). In brief, the transfected cells were seeded in 96-well plates at a density of 3000 cells/well in 100 µL at 37℃. At the three evaluation time points above, the plates were incubated in the dark at 37℃ for 2 h after adding 10 µL CCK8 solution. Spectrometric absorbance of each well at 450 nm was determined using SpectraMax M5 microplate reader (Molecular Devices). Wells with no cells were used as the control.

A scratch-wound healing assay was implemented as previously reported^159^. Briefly, the transfected cells were cultured in a 6-well plate with 2 mL DMEM supplemented with 10% FBS per well. After cells reached confluency, a vertical scratch was created in the cell monolayer using a sterile pipette tip. The wells were then washed with PBS, replenished with DMEM medium, and photographed with a light microscope (Olympus). The scratch width was estimated using ImageJ, and migration rate of wound healing was calculated according to the following formula^160^: *Migration rate*= 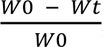, where W_0_ and W_t_ indicate initial wound width at 0 h and t h after wounding, respectively.

#### F-actin staining

For visualization of the actin cytoskeleton, the A7r5 cells were fixed with 4% paraformaldehyde (Solarbio) for 20 min, at 24 h, 48 h, and 72 h after transfection. F-actin staining was performed with Alexa-Fluor 488 phalloidin (Thermo Fisher Scientific) in the dark at room temperature for 30 min. The nuclei were counterstained with DAPI (Invitrogen) at room temperature for 5 min. Images were captured with a fluorescence microscopy (Olympus).

#### Generation of *Luc7l* knock-out mice

The *Luc7l* knock-out mice were generated by co-microinjection of Cas9 mRNA and gRNA into the cytoplasm of C57BL/6 zygotes (Extended Data Fig. 13a). Founders with frameshift mutations were screened with PCR (Supplementary Table 16 and Extended Data Fig. 13b). The founders with positive knock-out results were intercrossed to generate homologues *Luc7l*^−/−^ mice. Two guide RNAs (gRNAs) (gRNA1: 5’-CAGTTTCTTAGTGTATATGA-3’; gRNA2: 5’-CACATGAATGAATGTAGTTG-3’) were designed targeting exon 2 of the Luc7l-202 transcript (ENSMUST0000010014976.9) using the online CRISPR tool (http://crispr.mit.edu/). All animals were housed under the conditions with a 12 h light/dark cycle and free access to food and water.

#### *In vivo* fluorescence imaging

Eight-week-old wild-type (WT) and *Luc7l*^−/−^ mice were fasted for 12 h before imaging experiment. Then the fasting mice were orally gavaged with 4-kDa FITC dextran (Sigma-Aldrich) at a dose of 600 mg/kg body weight (dissolved in PBS at 80 mg/mL). Subsequently, these mice were anesthetized with isoflurane (induction 5%, maintenance 2% isoflurane) and imaged using an AniView 100 live animal imaging system (Biolight Biotechnology Co. Guangzhou, China) with excitation at 465 nm and emission at 600 nm.

#### Gastric emptying evaluation

For gastric emptying assessment, we folllowed the established phenol red marker meal protocol^161,162^. After 24 h fasting (water ad libitum), WT and *Luc7l*^−/−^ mice received 0.5 mL phenol red solution (0.5 mg/mL in 1.5% methylcellulose) by oral gavage. At 20 min post-gavage, mice were euthanized via CO_2_ asphyxiation with cervical dislocation confirmation. The stomach was excised after dual ligation of cardiac and pyloric orifices. The stomach was cut into pieces and homogenized with its contents in 25 mL of 0.1 M NaOH. After trichloroacetic acid (TCA) precipitation and centrifugation (3,000*g*, 20 min, 4°C), supernatant absorbance was measured at 560 nm. The concentration of phenol red in the stomach representing the gastric emptying (GE) was calculated as *GE* (%) = 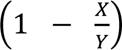 × 100, where X is absorbance of phenol red recovered from the stomachs of transgenic mice sacrificed 20 min after test meal, and Y is absorbance of phenol red recovered from the stomachs of wild-type mice (killed at 0 min following test meal).

## ACKNOWLEDGEMENTS

This study was financially supported by grants from the National Key Research and Development Program of China (Nos. 2021YFD1200900, 2021YFF1000703, 2022YFE0113300, 2022YFF1003402, and 2021YFD1300904) and the National Natural Science Foundation of China (Nos. 32272845 and 32320103006), the Project of Northern Agriculture and Livestock Husbandry Technical Innovation Center, Chinese Academy of Agricultural Sciences (BFGJ2022002) and the Biological Breeding-National Science and Technology Major Project (2023ZD0407106). We also thank the High-performance Computing Platform of China Agricultural University. We thank associate Prof. Yanling Ren from the Shandong Binzhou Animal Science and Veterinary Medicine Academy and Prof. Yonggang Liu from Yunnan Agricultural University for their support in sampling.

## AUTHOR CONTRIBUTION

M.H.L. conceived and designed the study. M.H.L. supervised and managed the project with contributions of S.S.X., and S.G.J. Q.X.H., S.S.X., W.T.W., and Y.H.Z. performed the bioinformatic analyses, and drafted the initial manuscript. Q.X.H., W.T.W., Y.H.Z., J.H.H., and S.S.X. participated in sample collection and resource generation. J.X.Z. contributed to the cell culture. J.H.H. contributed to the generation of overexpression mice. M.H.L., S.S.X. and S.G.J. interpreted the analytical results, wrote and revised the manuscript. All the authors reviewed, edited and approved the final manuscript.

## Data and code availability

All raw data analyzed in this study are publicly available for download without restriction from the Sequence Read Archive (SRA) database in NCBI under accession numbers PRJNA1369768. The mass spectrometry proteomics data have been deposited to the ProteomeXchange Consortium (https://proteomecentral.proteomexchange.org) via the iProX partner repository with the dataset identifier PXD073673.The codes used in this study are available at the GitHub repository: https://github.com/qixuan961017-apple/stomach.

## Competing Interests

The authors declare no competing interests.

## Notes

### Competing Interest Statement

The authors have declared no competing interest.

